# A brain locus for viable gestation

**DOI:** 10.64898/2026.09.15.751878

**Authors:** Sakura Tanaka, Joseph R. Knoedler, Vinicius Miessler de Andrade Carvalho, Maricruz Alvarado, Yichao Wei, Renzhi Yang, Adarsh Tantry, Sheruni A. E. Pilapitiya, Bibudha Parasar, Miao Wang, Chung-ha O. Davis, Lele Cui, Mariko H. Foecke, Diana J. Laird, Longzhi Tan, Nirao M. Shah

## Abstract

The expectant mother’s body is systemically remodeled by her hormones to weather the physiological vicissitudes of pregnancy. In contrast to other reproductive organs, how the expectant mother’s brain promotes these system-wide changes is largely unexplored^1,2^. Here we show that pregnancy induces profound changes in gene expression and identity of hormone-sensitive neurons and report a previously unknown class of such cells that is required for viable pregnancy. We performed RNA sequencing from late mid-gestation mice of four estrogen receptor alpha-expressing populations from hypothalamus and amygdala that regulate reproductive behaviors altered during gestation. Each of these populations undergoes such large, specific transcriptional shifts during pregnancy that these changes even exceed their transcriptional differences between the sexes^3^. The gene expression changes during pregnancy also imbue particular transcriptomically-defined neuronal types within these four populations with new molecular identities. Targeted ablation of one such neuronal type, POA^Npy2r^ cells, precludes implantation and abrogates viable pregnancy. Taken together, we have uncovered an essential role of hormone-sensitive neurons in the brain in sustaining pregnancy. The etiology of spontaneous gestational loss, which afflicts ∼15% of human pregnancies^4–6^, remains idiopathic in many cases, and our findings suggest a brain-based mechanism that contributes to such events. More broadly, we provide a molecular and cellular foundation to study gestational processes in health and disease from the perspective of the pregnant brain-body axis.

---

Pregnancy imposes a physiological burden on the expectant mother, requiring adaptations across virtually every organ system^1,2^. Such adaptations are, in large part, orchestrated by the hormonal *milieu* of gestation, and they result in many obvious changes. These include changes in ovulation, appetite, body weight, sexual behavior, parenting, and cognitive functions such as mood and odor perception. Together, this systemic remodeling ensures health of the expectant mother and growing baby.

How the pregnant brain detects and governs these body-wide changes during gestation is poorly understood. Moreover, and in contrast to prior studies pinpointing neuronal pathways that control ovulation, mating, or maternal behaviors, how the brain regulates pregnancy itself remains unclear. We reasoned that, similar to the rest of the body, the hormonal shifts of gestation act on target neurons in the brain to sustain pregnancy. Four populations of estrogen receptor alpha (ERα or Esr1)-expressing neurons in the female amygdala and hypothalamus exert inordinate influence on mating, parenting, and ovulation^3,7–21^: the bed nucleus of the stria terminalis principal nucleus (BNSTpr), medial amygdala (MeA), preoptic hypothalamus (POA), and ventromedial hypothalamus ventrolateral sector (VMHvl). Each of mating, parenting, and ovulation is markedly altered during gestation, suggesting that these centers are important targets of pregnancy-related hormones. Indeed, these neurons also co-express the receptor for progesterone (PR), a critical hormone for gestation, and they undergo extensive transcriptional and cellular remodeling across the mouse estrous cycle^3,22^. We therefore asked if these Esr1+ populations are also reconfigured during pregnancy.

We used genetically-targeted RNA sequencing to sensitively capture potential transcriptional changes and cell identity transformations of Esr1+ neurons in the BNSTpr, MeA, POA, and VMHvl during late mid-gestation. We found that gestation induced profound alterations in gene expression that were unique to each of these four populations, indicating specific rather than general neuronal adaptations to the various demands of pregnancy. These transcriptional changes were substantive enough to promote shifts in molecular identities of particular transcriptomically-defined neuronal types within these four Esr1+ populations. Targeted ablation of one such neuronal type, POA^Npy2r^ cells, abolished viable pregnancy, revealing a causal role for this neuronal type in sustaining gestation. Together, we have found that gestation reconfigures the molecular and cellular landscape of the brain of the expectant mother in a manner that is both region-specific and functionally essential for a viable pregnancy.

## Transcriptional landscape of the pregnant brain

In principle, pregnancy hormones could induce shared or unique transcriptional adaptations across different neuronal populations to meet the demands of gestation. Shared transcriptional adaptations would suggest a generic response to pregnancy by neuronal populations otherwise preconfigured to support gestation. Unique adaptations on the other hand would indicate that, regardless of their lineage history, different neuronal populations need to mount distinct gene expression programs to support pregnancy. To distinguish between these possibilities, we utilized a high throughput approach that affords deep and reliable coverage of the transcriptome. We performed translating ribosome-affinity purification followed by deep sequencing (TRAPseq^3,23^) of the four Esr1+ populations from pregnant (gestational day 14, G14) *Esr1^Cre^;RiboTag* mice to discern global alterations of the transcriptome in these cells^23,24^ (Fig. 1a and Extended Data Fig. 1a-e). We compared transcriptional profiles from pregnant females (F_P_) to those we had collected from virgin females hormonally induced to be in estrus (sexually receptive, F_R_) or diestrus (sexually unreceptive, F_NR_)^3^. We found 2,788 differentially expressed genes in pregnancy (pDEGs) across these four populations (≥1.5-fold change cutoff; 1.9 median fold change), with similar numbers of pDEGs between F_P_ and F_R_ (1,425) and F_P_ and F_NR_ (1,363) comparisons (Fig. 1b, Extended Data Fig. 2a, and Supplementary Tables 1, 2). These pDEGs mapped to 1,709 coding genes – or ∼8% of coding genes – that are distributed across all chromosomes, and complementary validation with *in situ* hybridization confirmed differential expression of all 25 pDEGs tested (Fig. 1c and Extended Data Fig. 1f, 2b).

**Figure 1:**
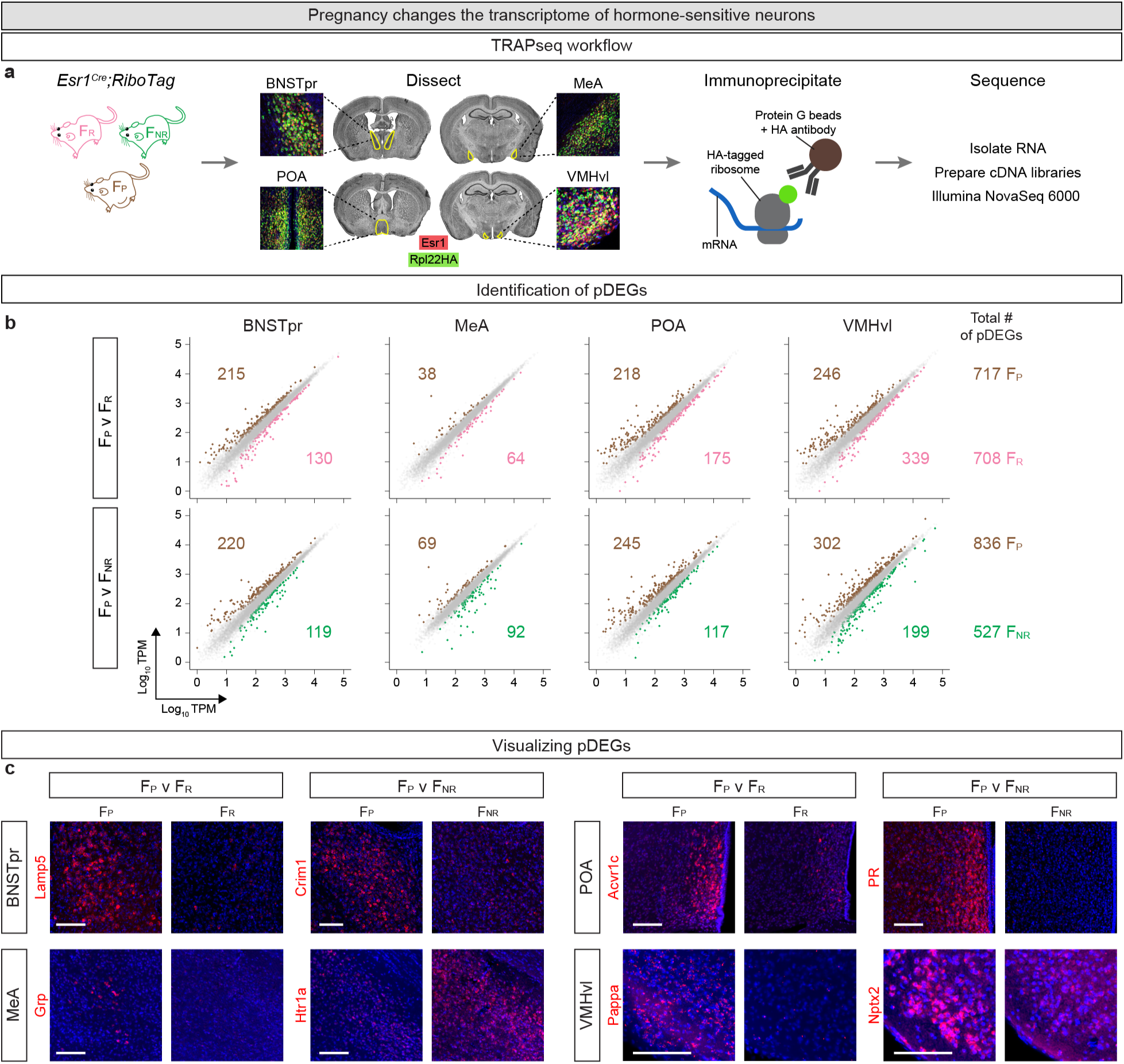
TRAPseq identification of pDEGs. **a.** TRAPseq workflow. **b.** Scatter plots of pDEGs in different Esr1+ populations. Dots represent DEGs with >1.5-fold change and FDR-adjusted *p* value <0.05 (colored) or genes that did not meet both criteria (gray). Colored numbers enumerate pDEGs upregulated in that condition and comparison for each region or (far right) for all regions combined. TRAPseq identified 1,425 and 1,363 pDEGs in F_P_ v F_R_ and F_P_ v F_NR_ comparisons. v, versus; TPM, transcripts/million; *N* = 3/condition. **c.** Representative ISH images for pDEGs. ISHs confirm TRAPseq data showing higher expression of *Lamp5*, *Grp*, *Acvr1c*, and *Pappa* in F_P_ compared to F_R_; *Crim1*, *PR* and *Nptx2* in F_P_ compared to F_NR_; and *Htr1a* in F_NR_ compared to F_P_. Sections in coronal plane, with medial to the right and dorsal at the top of the panels, for all Figures. Red, mRNA; blue, DAPI. Scale bars = 100 µm. *N* = 2/condition/gene.

We tested whether pregnancy induced shared or unique transcriptional changes in relation to different physiological states (F_R_ and F_NR_) and brain regions. Most pDEGs (≥59%) were in fact restricted to individual pairwise comparisons within each Esr1+ population (Fig. 2a). Moreover, principal component analysis (PCA) showed that F_P_, F_R_, and F_NR_ occupied distinct states in PC space – with F_P_ clearly separated from F_R_ and F_NR_ in PC1 – indicating that pregnancy is a discrete state rather than one on a continuum with virginal states (Extended Data Fig. 3a). Unsupervised segregation of pDEG patterns using *k*-means clustering also revealed differential distribution of pDEGs across each physiological state that was robust to pooling across populations (Extended Data Fig. 3b-c). Most pDEGs were also unique to each brain region (≤25% common to ≥2 populations) such that each *k*-cluster also comprised unique sets of pDEGs and PCA showed that pDEGs segregated by brain region rather than physiological states (Fig. 2b, Extended Data Fig. 3d and Supplementary Table 2). Overall, we only identified 3 pDEGs that were similarly changed in all four populations between pregnancy and virginal states. *Cacng8*, encoding a subunit of a calcium channel and *Hs3st4*, encoding a sulfotransferase for heparan sulfate, were upregulated in gestation, whereas *Rbm3*, encoding an RNA-binding protein implicated in ovarian cancer^25^, was downregulated in pregnancy. Intriguingly, *Galanin* is upregulated in POA and BNSTpr and downregulated in MeA and VMHvl. Galanin+ POA neurons regulate parenting, and our findings at G14 raise the possibility that the transcriptional changes in these cells portend maternal behavior even earlier than previously described^8,15,26^. Regardless, we did not identify a large set of pDEGs that paints a shared molecular representation of pregnancy. Together, pregnancy induces transcriptional adaptations that are distinct for different populations and reproductive states.

**Figure 2:**
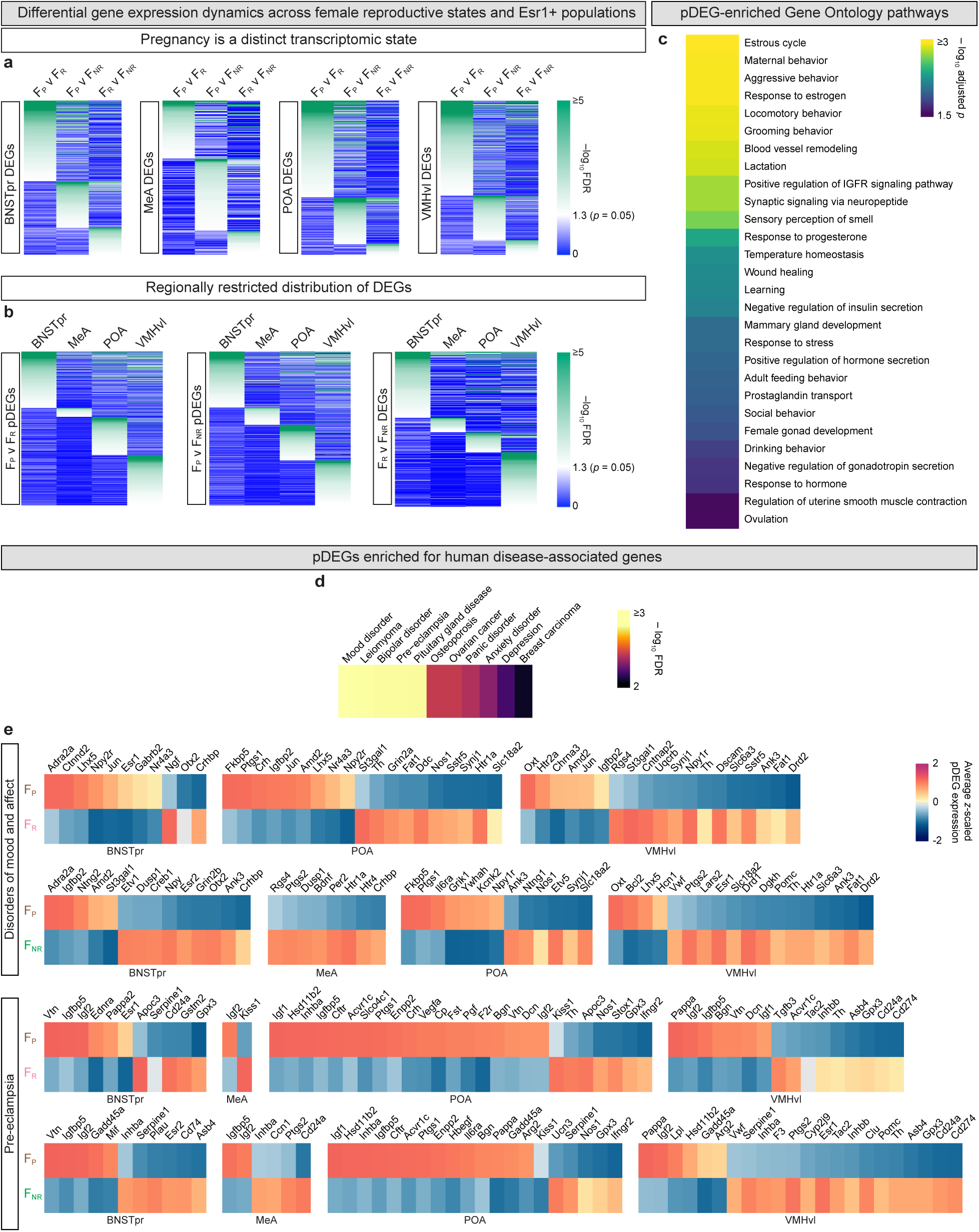
Differential gene expression across reproductive states and Esr1+ populations. **a.** Heatmap of FDR-adjusted *p* values of individual DEGs for the relevant pairwise comparison shows that most DEGs within an Esr1+ population are only differentially expressed between one pair of female reproductive states. **b.** Heatmap of FDR-adjusted *p* values of individual DEGs for the relevant pairwise comparisons illustrating that most DEGs are restricted to one Esr1+ population. Heatmap of F_R_ and F_NR_ comparison is adapted from a previous study^3^. **c.** GO Biological Process pathways enriched among the combined set of pDEGs (merged from all Esr1+ populations). Significantly enriched GO pathways (adjusted p-value < 0.05) related specifically to female biology or pregnancy are shown. **d.** Human diseases significantly associated with the combined set of pDEGs (merged from all Esr1+ populations). Significantly associated diseases (FDR-adjusted p-value < 0.05) related specifically to female biology or pregnancy are shown. **e.** Heatmap of expression of pDEGs associated with disorders of mood and affect (top 2 rows) and pre-eclampsia (bottom 2 rows). Gene-disease associations were obtained from the Disease Ontology database. Scale = *z*-scored expression of pDEGs centered at zero for comparisons between pregnant and virgin states.

We queried pDEGs for associations with biological processes and human disease. Gene Ontology analysis^27,28^ showed enrichment for pDEGs in diverse signaling pathways, including most intriguingly in pathways related to estrous cycle (including *Cckbr, Esr2, PR*)^29–32^, female reproductive organs (including *Esr1, Inhba, Oxtr, Wnt4*)^33–37^, nursing (including *Avpr1a, Ccnd1, Oxtr, Stat5a*)^38–41^, and bone remodeling (including *Ccn1, Bmp6, Acvr2b*)^42–44^ (Fig. 2c and Supplementary Table 3). These findings suggest shared gene expression programs between the brain and peripheral tissues that undergo major adaptations during pregnancy. In agreement with this notion, querying the Disease Ontology database^45^ showed enrichment of pDEGs for neuro-psychiatric disorders that are more common in pregnancy as well as pre-eclampsia (Fig. 2d-e) and diseases of female reproductive organs (including *Brca1, Esr1, PR, Rbm3*)^25,46–48^ (Fig. 2d and Supplementary Table 3).

Together, we find that the brain adapts to pregnancy via a genome-wide, gestation-specific transcriptional program that is distinct for different sex hormone-responsive neuronal populations. In addition, this transcriptional program shares features with non-neural gene expression networks exclusive to women in health and disease.

## Cellular landscape of the pregnant brain

Given the transcriptomic heterogeneity of neurons in each of the four Esr1+ populations^3,22,49–52^, we mapped the distribution of pDEGs at cellular resolution with single-nucleus RNA sequencing (snRNAseq) of all four Esr1+ populations from G14 *Esr1^Cre^;SunTag* mice^53^ (Fig. 3a and Supplementary Table 1). In total, we sequenced 24,849 high quality Esr1+ nuclei across all four regions. Unsupervised clustering of all libraries (F_P_ and F_R_ and F_NR_^3^) showed that Esr1+ nuclei co-embedded in UMAP (Uniform Manifold Approximation and Projection) space by region of origin, indicative of minimal batch-effects from sequencing (Extended Data Fig. 4a). Data from F_P_ neurons integrated with our previously published taxonomy of Esr1+ neuronal types in virgin F_R_ and F_NR_ mice from these brain regions^3^, as determined by unbiased direct transcriptome comparisons between neuronal types (Fig. 3b and Extended Data Fig. 4b-d). This co-embedding revealed an additional 14 neuronal types present in all physiological states that were a consequence of adding data from more neurons to re-build the taxonomy (Extended Data Fig. 5a-d and Supplementary Table 4). We also observed differential enrichment of 15 neuronal types across all four Esr1+ populations in one or two physiological states (Extended Data Fig. 5e). Taken together, the four Esr1+ populations we have sequenced from F_R_, F_NR_, and F_P_ mice comprise 154 transcriptomically-defined neuronal types, each of which expresses type-specific marker genes (Extended Data Fig. 5a-d and Supplementary Table 4).

**Figure 3:**
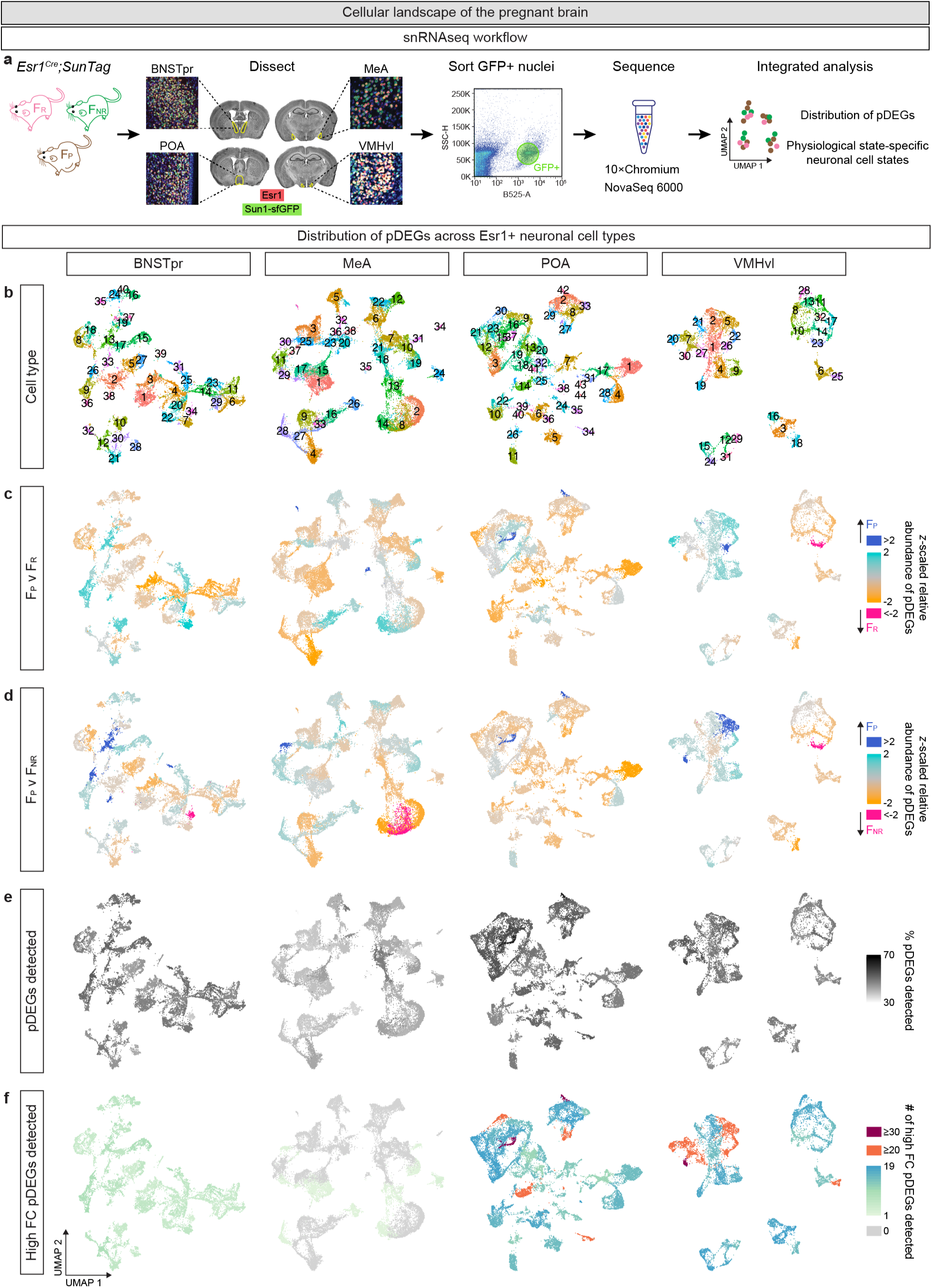
Pregnancy in Esr1+ populations at single-cell resolution. **a.** snRNAseq workflow. **b.** UMAP representation of Esr1+ neuronal types. **c-d.** Neuronal types colored based on *z*-scaled relative abundance of pDEGs, indicating whether they exhibit a bias towards upregulated (turquoise/blue) or downregulated (orange/pink) expression in F_P_ compared to F_R_ (**c**) or F_NR_ (**d**). **e.** UMAP shows that each neuronal type expresses ≥30% of pDEGs in all Esr1+ populations. **f.** UMAP shows distribution of high fold-change pDEGs (>4-fold difference from F_R_ and/or F_NR_), showing more restricted distribution across Esr1+ neuronal types than that for the full complement of pDEGs. FC, fold change.

We next examined gene expression patterns and other features of these 154 transcriptomically-defined neuronal types. We found pDEGs in all neuronal types, including excitatory and inhibitory types in each brain region, regardless of the comparison to F_R_ or F_NR_ states (Fig. 3c-d and Extended Data Fig. 5). Moreover, any individual neuronal type expressed ≥30% of pDEGs within the corresponding Esr1+ population (Fig. 3e). Despite the ubiquity of pDEGs across all transcriptomically-defined neuronal types, a small subset was significantly enriched or depleted in a few Esr1+ neuronal types (Extended Data Fig. 4e). With few exceptions, we found comparable abundance of pDEGs in Esr1+ neuronal types within each region, ranging from a median of 50 to 263 per neuronal type in the MeA and VMHvl, respectively (Extended Data Fig. 4e and 6a). Such uniformity of pDEG distribution within individual regions belies the unique set of pDEGs detected between any pair of cell types (Extended Data Fig. 6b).

A surprising number of pDEGs (363, comprising 13% of all pDEGs, and encoded by 241 genes) exhibited a large (>4) fold change regardless of the comparison to F_R_ and F_NR_ states. Analysis of these high-fold change pDEGs showed associations in Gene and Disease Ontology databases that were similar to the entire collection of pDEGs (Extended Data Fig. 6c-d). However, and in contrast to the uniform distribution of all pDEGs across brain regions and neuronal cell types, high-fold change pDEGs were selectively enriched in neuronal types in the BNSTpr, POA, and VMHvl compared to the MeA (Fig. 3f). Indeed, 27/38 MeA neuronal types did not express such pDEGs. Many neuronal types in the POA (6) and VMHvl (8) expressed >20 high-fold change pDEGs, including VMHvl^Cckar^(F_R_-specific and essential for female sexual behavior^3,18^) and POA^Npy2r^ and POA^Svil^ (both F_P_-specific; Extended Data Fig. 5e). Nevertheless, the observation that neuronal types present in all three physiological states expressed high fold-change pDEGs suggests functional specialization even within these shared clusters of neurons. Together, pregnancy elicits unique transcriptional changes in all transcriptomically-defined neuronal types within the four Esr1+ populations. Moreover, these gene expression changes are large enough to transform some neuronal types into distinct cell states in virgin and pregnant mice.

## Regulatory landscape of the pregnant brain

We found dynamic changes in the expression of 143 transcription factors (TFs) between pregnancy and virgin states across Esr1+ populations (Fig. 4a). These pDEG-encoded TFs (pDEG-TFs), similar to other pDEGs, were expressed in distinct patterns in different neuronal cell types across the four regions. This suggested the possibility that these pDEG-TFs regulated pregnancy-induced changes in gene expression. We used the SCENIC pipeline to determine whether these pDEG-TFs regulated expression of other pDEGs, thereby assembling into regulons comprising regulator pDEG-TFs associated with respective sets of regulated pDEGs^54^. In fact, a large fraction of pDEG-TFs (ranging from 41.7% in the POA to 73.3% in MeA) formed such regulons that in turn accounted for a majority of all pDEGs (Fig. 4b). pDEGs not currently members of pDEG-TFs based regulons may eventually turn out to be regulated by pDEG-TFs as SCENIC expands its coverage of the genomic regulatory space; alternatively, these pDEGs may be regulated by regulons containing a TF that is not a pDEG (Supplementary Table 5). In any event, we found that many pDEG-TFs (43) were predicted to regulate other pDEG-TFs (69), indicative of a nested regulon structure in pregnancy (Extended Data Fig. 7a). Notably, PR is a pDEG in the BNSTpr, POA, and VMHvl, and it forms regulons comprising pDEGs as well as other pDEG-TFs (Supplementary Table 5), in agreement with the prominent role of progesterone in sustaining pregnancy. The POA^Npy2r^ (#37) and POA^Svil^ (#42) neuronal types which show the largest transcriptional shifts in pregnancy also show enrichment and depletion of multiple pDEG-TFs, including PR (Fig. 4a). Our findings suggest that the transcriptomic changes of pregnancy arise from, at least in part, pDEG-TFs that regulate expression of each other and other pDEGs. In principle, the wholesale changes in gene expression during pregnancy could have arisen from changes in genome architecture. However, probing a well-characterized and validated dimorphic cell type revealed no changes in chromatin contacts during pregnancy^3,18,55–57^ (Extended Data Fig. 7b-e and Supplementary Table 1). Taken together, our findings indicate that the large changes in gene expression during pregnancy emerge from dynamic changes in the regulatory network of TFs in Esr1+ neurons.

**Figure 4:**
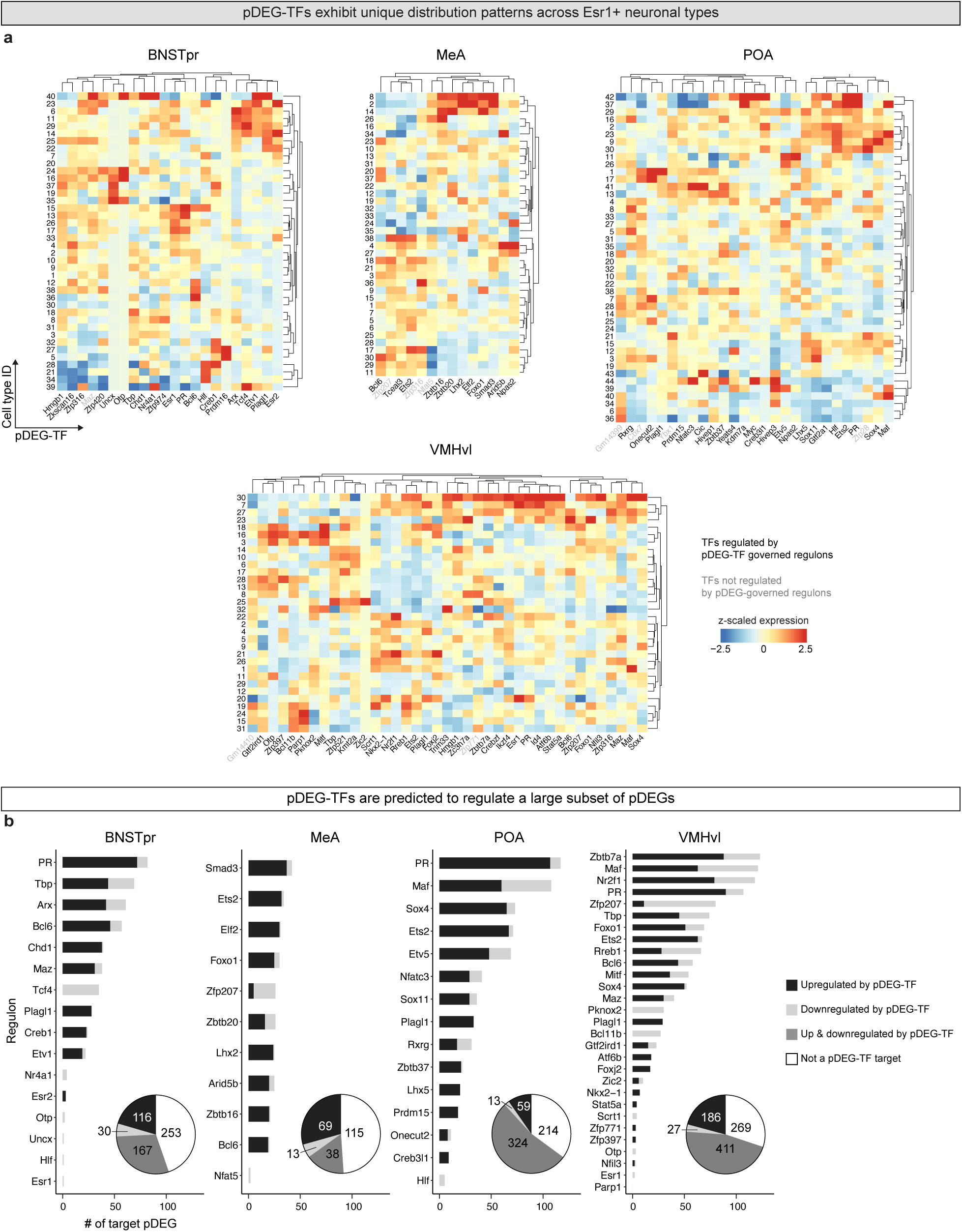
Regulatory logic of gene expression changes in pregnancy. **a.** pDEG-TFs exhibit a unique expression pattern across cell types in each Esr1+ population. Most pDEG-TFs are contained within pDEG-TF governed regulons. Rows and columns are hierarchically clustered using Euclidean distance. **b.** More than 50% of pDEGs are predicted to be regulated by pDEG-TFs in all Esr1+ populations.

## Viable pregnancy requires POA^Npy2r^ neurons

We tested the functional relevance of the GABAergic POA^Npy2r^ neuronal type during gestation. We chose to target these neurons because they exhibit the largest transcriptional shift in pregnancy, express >30 high fold-change pDEGs, and can be genetically targeted using the *Npy2r^Cre^*mouse strain^58^ (Fig. 3c-f and Extended Data Fig. 6a). This cell type is exclusive to pregnancy (#37), and it likely results from a transcriptional shift of cell type #23 in F_R_ and F_NR_ states, as evidenced by shared marker genes and a corresponding depletion of #23 in F_P_ (Fig. 5a-b and Extended Data Fig. 5e). Further impetus to functionally interrogate POA^Npy2r^ neurons was provided by our observation that the human female POA also contains a circumscribed cluster of neurons expressing NPY2R within a larger GABAergic population^59^ (Fig. 5c).

**Figure 5:**
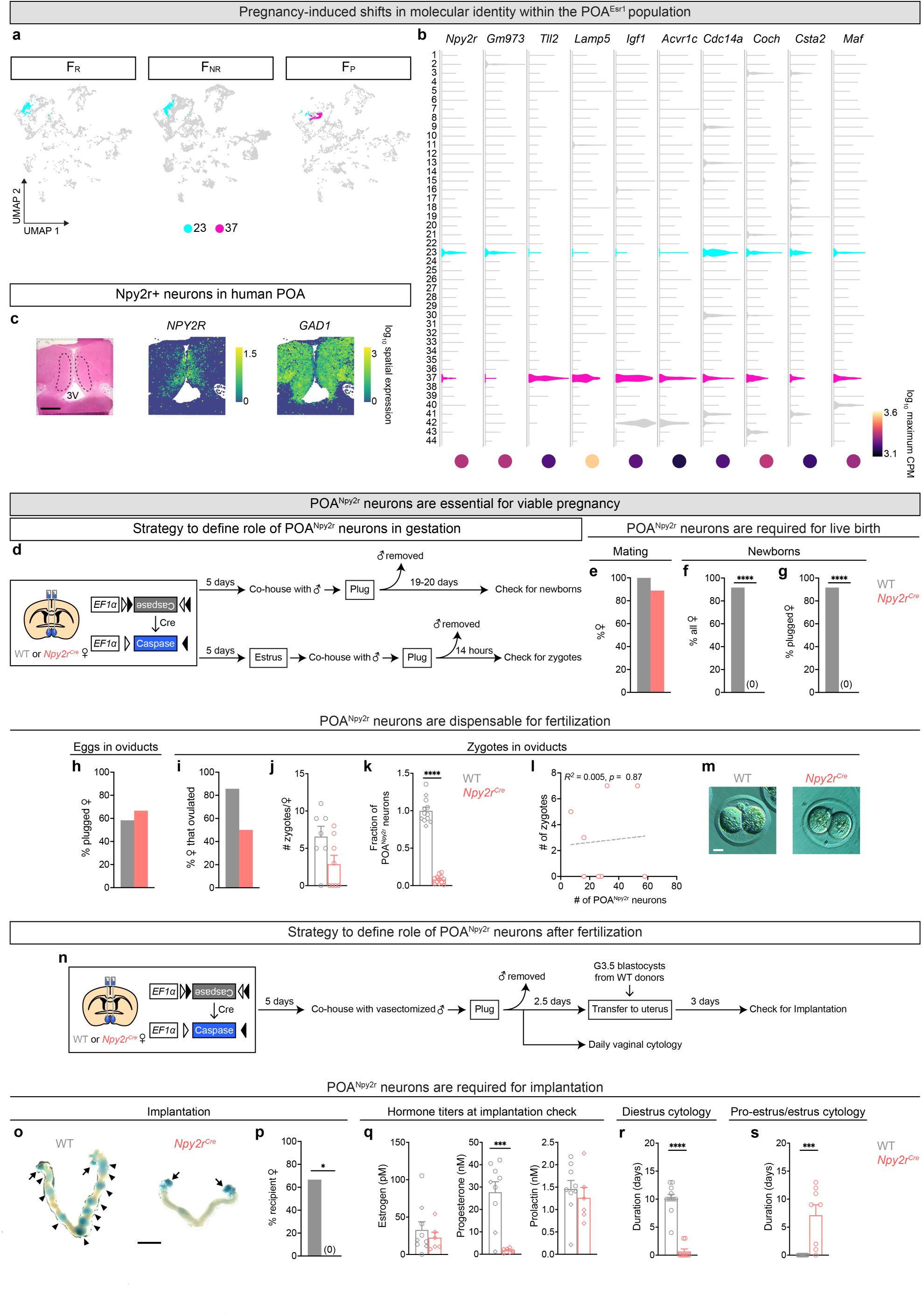
POA^Npy2r^ neurons are required for viable pregnancy. **a.** UMAP representation of female POA^Esr^^1^ neurons shows that neuronal type #37 is restricted to the F_P_ state. **b.** Neuronal type #37 in F_P_ state expresses marker genes and pDEGs shared with neuronal type #23 in F_R_ and F_NR_ states. **c.** NPY2R and GAD1 are expressed in adult human female POA. Panel on left is POA stained with hematoxylin-eosin. Areas demarcated with dashed lines correspond to the POA. 3V, 3^rd^ ventricle; scale bar = 3 mm. **d.** Workflow to define role of POA^Npy2r^ neurons in gestation. **e.** No difference between females ± POA^Npy2r^ neurons in successful copulation (documented with vaginal plug). **f-g**. Absence of newborns following ablation of POA^Npy2r^ neurons. **h-m**. Comparable ovulation (eggs observed in oviducts; **h**), fertilization (zygotes observed in oviducts that developed into 2-cell embryos *in vitro*; **i, j**) in females ± POA^Npy2r^ neurons. POA^Npy2r^ neuron number plotted as fraction of neurons compared to control following ablation of these neurons (**k**). No correlation between extent of POA^Npy2r^ neuronal loss (number of surviving neurons plotted) and number of zygotes (**l**). Two-cell embryos *in vitro* in females ± POA^Npy2r^ neurons (**m)**. Circles and diamonds, females with and without eggs, respectively; scale bar = 25µm. **n**. Workflow to define role of POA^Npy2r^ neurons post-fertilization. **o-p**. Embryo implantation sites stained with Chicago Blue (blue bulges in uterus, black arrowheads) are present in a majority of control (WT) females and absent in those lacking POA^Npy2r^ neurons. Ovaries stained blue (black arrows) are visible in both groups of females. Uterus is elongated and dilated in control females, as expected for early pregnancy. Scale bar = 5 mm. **q.** Females lacking POA^Npy2r^ neurons have low progesterone at the time of implantation check. Circles and diamonds, females with and without implantation sites, respectively. **r-s.** Ablation of POA^Npy2r^ neurons induces prolonged proestrus or estrus vaginal cytology following successful copulation in contrast to the prolonged diestrus of control females. Sample sizes: WT*, N* = 12 and *Npy2r^Cre^*, *N* = 10 (**e-g**); WT, *N* = 12 and *Npy2r^Cre^*, *N* = 12 (**h-m**); WT, *N* = 9 and *Npy2r^Cre^*, *N* = 6 (**o-q**); WT, *N* = 11 and *Npy2r^Cre^*, *N* = 8 (**r-s**). Statistical analyses: Fisher’s exact test (**e-i**, **p**); Student *t*-test (**k, q** center and right panels); Mann-Whitney test (**j**, **q** left panel, **r-s**); Pearson’s correlation coefficient (**l**). *, *p* < 0.05; ***, *p* < 0.001; ****, *p* < 0.0001.

The onset of any potential gestagenic role of POA^Npy2r^ cells is unknown. We reasoned that, analogous to a genetic loss-of-function approach, ablating this neuronal type would reveal any essential function in pregnancy in an unbiased manner. Indeed, cellular loss or damage has provided deep insights on neuronal contributions to physiology and behavior^60,61^. Accordingly, we ablated POA^Npy2r^ neurons by delivering a virally-encoded Cre-dependent, cell-autonomously lethal toxin caspase-3 to the POA of *Npy2r^Cre^* or WT (wildtype, control) females, co-housed them with a WT male, and checked for successful copulation (evidenced by an ejaculatory vaginal plug) every day^9^ (Fig. 5d and Extended Data Fig. 8a-b). We observed no difference in mating success between experimental and control females, both in terms of the latency to plug and percent females plugged (Fig. 5e and Extended Data Fig. 8c). However, and in contrast to the majority (>90%) of control females, none of the experimental females delivered pups (Fig. 5f-g), revealing an absolute requirement of POA^Npy2r^ neurons for viable pregnancy.

Mice start gaining weight after G10 but none of the females in whom we had ablated POA^Npy2r^ neurons displayed discernible weight gain, indicative of fetal loss prior to G10 (Extended Data Fig. 8d). Given the absence of embryos this early in pregnancy, we tested if ablation of POA^Npy2r^ neurons even precluded fertilization. Fertilization occurs rapidly after plug formation in mice^62^ and we therefore modified our experimental approach following delivery of Cre-dependent caspase-3 to the POA of *Npy2r^Cre^* females (Fig. 5d). We co-housed estrus females with WT males, checked for plugs every 6 hours, and examined oviducts for eggs 14 hours after observing a plug, when we could reliably observe single-cell embryos (zygotes) in WT females. We observed no discernible difference in the number of females with eggs or zygotes in the oviduct or the number of zygotes between experimental and control mice following successful copulation (Fig. 5h-m and Extended Data Fig. 8e). Moreover, ovaries from these two cohorts of mice contained comparable follicles and corpora lutea in different stages, confirming that they had successfully ovulated (Extended Data Fig. 8f-g). In agreement with the comparable fertilization indices between females ± POA^Npy2r^ neurons, we detected no difference in serum titers of key hormones, including estrogen, progesterone, and gonadotropins (Extended Data Fig. 8h). Importantly, serum prolactin levels rise rapidly after mating and are critical to sustain gestation^63–66^, and we found no difference in titers of this hormone between experimental and controls (Extended Data Fig. 8h). Females lacking POA^Npy2r^ neurons also appeared comparably healthy, with a typical groomed appearance and similar levels of blood glucose (Extended Data Fig. 8h). Together, these findings demonstrate that POA^Npy2r^ neurons regulate events downstream of fertilization to enable viable gestation.

A central event following fertilization is implantation, which occurs at ∼G4 in mice and allows the growing embryos to invade the uterus and obtain sustenance from the mother^62^. We therefore tested if females lacking POA^Npy2r^ neurons were able to implant embryos. We employed the standard protocol for mouse transgenesis to transfer embryos from WT donor mice into experimental females lacking POA^Npy2r^ neurons^67,68^ (Fig. 5n). Three days after receiving WT blastocysts, we observed numerous implant sites^69^ in WT (control) mice but none in females lacking POA^Npy2r^ neurons (Fig. 5o-p and Extended Data Fig. 8i-k). Accordingly, ablation of these cells in a separate set of WT-blastocyst recipients abolished viable gestation such that these females neither gained weight nor birthed a litter when allowed to go to term (Extended Data Fig. 8m-q). Together, our findings show that POA^Npy2r^ neurons are essential for implantation.

Females ± POA^Npy2r^ neurons did not differ in levels of estrogen, prolactin, gonadotropins, and glucose despite the absence of implanted embryos in *Npy2r^Cre^* females (Fig. 5o-q and Extended Data Fig. 8l). Nevertheless, mice lacking POA^Npy2r^ neurons did not display the increased progesterone titers that accompany pregnancy^70^ (Fig. 5q). In addition to a rise in circulating progesterone and prolactin, successful copulation is followed by an exit from proestrus or estrus vaginal cytology to diestrus cytology that persists through pregnancy, cessation of ovulation, and perdurance of corpora lutea from the ovulation that preceded or accompanied mating^1,19,63,64,70–73^. These changes are also initiated in females who have successfully copulated with a vasectomized male in which case they are collectively referred to as pseudopregnancy, a state instrumental to enabling transgenesis in mice^68,71,73,74^. Within the context of normal serum prolactin and lowered progesterone (Fig. 5q), females lacking POA^Npy2r^ neurons lacked discernible corpora lutea (Extended Data Fig. 8r-s), indicative of cessation of new ovulatory cycles after mating. Moreover, ablation of POA^Npy2r^ precluded mated females from entering a physiological state with diestrus cytology such that they continued to show proestrus or estrus cytology (Fig. 5r-s and Extended Data Fig. 8t). Taken together, our findings show that POA^Npy2r^ neurons are essential to induce key physiological changes at implantation to sustain viable pregnancy.

## Discussion

The physiology and behavior of the expectant mother undergo a radical transformation to support her health and that of the growing embryo. The brain is the proximate arbiter of these changes and accordingly we demonstrate significant remodeling of gene expression and cellular states in the sex hormone-responsive neuronal populations we sampled. Indeed, the transcriptional shift during pregnancy exceeds the transcriptional differences between the sexes by 550 genes (2.5% of coding genes)^3^, underscoring the unique physiology and behavior of this lifecycle stage. These transcriptional shifts also lead to particular neuronal types changing their molecular identities during pregnancy. These changes are significant because ablation of one such neuronal type, which we estimate as comprising fewer than 0.001% of cells in the brain, abrogates viable gestation.

Our current findings and previous work on gene expression changes across the estrous cycle reveal that distinct female reproductive states – pregnancy, estrus, and diestrus – are characterized by equally distinct gene expression programs in the brain. We anticipate that similarly singular gene expression programs accompany lactation and menopause, which are two other major lifecycle events restricted to females. Such dynamic and stage-specific gene expression changes in adult females likely reflect evolutionary adaptations that enhance reproductive success.

Most pDEGs are unique to the Esr1+ populations we have examined in the BNSTpr, MeA, POA, and VMHvl. The regionally-restricted expression of most pDEGs is regulated by equally circumscribed layered transcriptional networks that are themselves regulated by pregnancy. There are many other sex hormone-responsive neuronal populations in the brain, and we expect that these cells will express an equivalently unique gene set during pregnancy. Such neurons may also cell non-autonomously modulate the transcriptional profiles of neighboring or synaptically connected neurons that are not responsive to sex hormones. In agreement with prior work^75,76^, we therefore expect that pregnancy elicits wholesale changes in gene expression across the brain.

Every transcriptomically-defined Esr1+ neuronal type we examined undergoes large changes in its gene expression profile. These ubiquitous transcriptional alterations may compensate for the physiological slings and arrows of pregnancy or they may promote changes in processes that sustain gestation. Together with our previous work^3^, we have now identified several female-specific transcriptional shifts in neuronal identities across estrus, diestrus, and pregnancy, indicative of the distinct demands of these different physiological states.

We interrogated the function of the neuronal type, POA^Npy2r^ cells, with the largest transcriptional shift in identity during pregnancy. Ablation of these neurons abrogated the ability of the uterus to implant embryos, resulting in failure of progression to term. *Npy2r* is expressed in POA^Npy2r^ cells in virgins prior to the transcriptional shift of pregnancy. These neurons may therefore already be primed to sustain gestation in virgins, or alternatively, rapidly undergo transcriptional changes to enable pregnancy following copulation. Regardless of the exact timepoint at which POA^Npy2r^ neurons support gestation, to our knowledge, we have identified the first transcriptomically-defined neuronal type that is essential for viable gestation. Prior studies have indicated that multiple brain regions, including the POA, regulate induction of pseudopregnancy^77,78^, primarily by assessing serum prolactin. We find that POA^Npy2r^ neurons regulate maternal changes in circulating progesterone and vaginal cytology after mating but not circulating prolactin or ovulation. Taken together with recent work^79^, our findings demonstrate that the many physiological changes that sustain early pregnancy are controlled in a modular manner by distinct sets of neurons.

An estimated 213 million women get pregnant every year globally^80^. Of these, ∼30 million pregnancies are thought to end in miscarriage (∼1 every second), including ones where gestational loss happens prior to self-knowledge or clinical confirmation of pregnancy^4–6^. The underlying etiology in many of these miscarriages remains unknown. We speculate that a subset of such events reflects failure of particular neural circuits to recognize, monitor, or adapt to gestation. The notion that neural circuits in pregnant women also adapt to and modulate gestation is consistent with imaging studies showing widespread alterations in their brains during healthy pregnancies^81–83^. Our analyses of the transcriptional changes of pregnancy also reveal genetic links to other, even more common afflictions of pregnancy, including those with a neural basis such as mood disorder as well as conditions thought to be peripherally based such as pre-eclampsia. Such associations during pregnancy could reflect previously unknown disease relevance of the neuronal populations we have queried or shared hormonally-regulated genetic networks that govern cellular function in the brain and elsewhere in the body. Our work now provides a new molecular and cellular framework to interrogate such shared as well as unique adaptations critical for gestation in the brain.

## Data availability

The sequencing data generated in this study have been deposited in the Gene Expression Omnibus (GEO) and are publicly available upon publication. Previously published sequencing data from F_R_ and F_NR_ mice can be accessed at GEO under accession numbers GSE183092 and GSE183093.

## Code availability

This paper does not report original code. All scripts used to generate the analysis and visualizations used in this manuscript are available from the lead contact upon reasonable request.

## Methods

### Animals

All animal procedures followed Stanford University Institutional Animal Care and Use Committee guidelines. *RiboTag, Esr1^Cre^,* and *SunTag* mice were purchased from Jackson Laboratories (JAX; 011029, 017913, 021039, respectively), *Npy2r^Cre^* mice (JAX; 029285) were generously provided by the Liberles lab (Harvard Medical School), and *Cckar^Cre^* mice (JAX; 037017) were previously generated in our lab^3,23,24,53,58^. Wild-type C57BL6/J females used for *in situ* hybridization, embryo transfer donors, and other experiments were purchased from JAX (000664) or bred in-house from JAX stock. Wild-type B6129SF1/J males used as sexually experienced males were also purchased from JAX (101043). Mice were group-housed by sex after weaning under a reversed 12:12 hour light:dark cycle (lights on at 1 AM) with food and water available *ad libitum*.

### Assessment and monitoring of pregnancy

To generate F_P_ mice for all studies, including TRAPseq and snRNAseq, females (8-10 weeks old) were co-housed with sexually experienced males and examined every morning for vaginal plugs. Females were weighed every other day between 9-11 AM starting at 5 days post-plug and every day at the same times starting at 10 days after co-housing. Given that not all females had confirmed plugs, and not all plugged females become pregnant, only females co-housed ≥14 days and showing ∼7 g (∼30% of starting body weight) weight gain were dissected. Females with fewer than 4 live embryos were discarded. Embryos from each female were enumerated and ≥2/female were assessed for Theiler stage^84^ for TRAPseq, snRNAseq, and Hi-C. Only females in which both embryos staged at TS21-24 were categorized as F_P_ mice and used for our studies.

### Dissection for TRAPseq and snRNAseq

F_P_ mice used for TRAPseq and snRNAseq were F1 *Esr1^Cre/+^*;*RiboTag/+* and *Esr1^Cre/+^*;*SunTag/+*, generated by mating *Esr1^Cre/Cre^* males with homozygous *SunTag* or *RiboTag* females, respectively. Homozygous parents were F1 progeny of heterozygous parents. Tissue preparation was conducted exactly as previously described^3^. In brief, females were deeply anesthetized with 2.5% Avertin and decapitated. Brains were sectioned coronally (500 µm) using a chilled brain matrix mold (BrainTree Scientific). Sections were floated in either chilled TRAPseq dissection buffer (100 mM KCl, 50 mM Tris-HCl pH 7.4, 12 mM MgCl_2_) or snRNAseq dissection buffer (0.25 M sucrose, 25 mM KCl, 5 mM MgCl_2_, and 20 mM Tricine-KOH pH 7.8). The BNSTpr, MeA, POA and VMHvl were identified using landmarks from a mouse brain atlas and dissected using a stereoscope^85^. Tissue was immediately frozen on dry ice and stored at -80°C until further processing (TRAPseq) or placed in cold homogenization buffer for subsequent nuclear extraction (snRNAseq). Each biological replicate comprised pooled tissue from 3 mice for TRAPseq and 5-7 mice for snRNAseq.

### TRAPseq

Translating ribosomes of Esr1+ cells from *Esr1^Cre^;RiboTag* females were purified exactly as previously described^3^. For each biological replicate from F_P_ mice, brain tissue from three dissected individual mice was pooled such that each replicate contained females with embryos that were comparably staged TS21-24. In brief, frozen brain tissue was thawed in a pestle homogenizer with 1 mL of ice-cold homogenization buffer containing 100 mM KCl, 50 mM Tris-HCL pH 7.4, 12 mM MgCL2, 1% IGEPAL, 1 mM dithiothreitol (DTT), 1 mg/mL heparin, 100 µg/mL cycloheximide, cOmplete mini protease inhibitor (Roche), and 40 U/µL murine RNAse inhibitor (NEB; M0314). The homogenate was clarified by centrifugation at 10,000 *g* for 10 minutes at 4°C and the post-mitochondrial supernatant was transferred to a new tube. 40 µL of the supernatant was mixed with 0.35 mL of RLT buffer (RNeasy Micro Kit; QIAgen; 74004) supplemented with 10 µL/mL β-mercaptoethanol (Input sample) and stored at -80°C overnight. For the IP sample, 2.5 µL of rabbit anti-HA antibody (Cell Signaling, 3724; RRID:AB_1549585) was added to the remaining homogenate and incubated in an end-over-end rotator at 4°C for four hours. The sample was then mixed with 50 µL of pre-washed paramagnetic protein G Dynabeads (Invitrogen) and incubated overnight at 4°C on an end-over-end rotator. After washing the beads with high-salt buffer (300 mM KCl, 50 mM Tris-HCl pH 7.4, 12 mM MgCl2, 1% IGEPAL, 1 mM DTT, 300 µg/mL cycloheximide), they were transferred to a new tube, collected with a magnet and the supernatant was removed. RLT buffer (0.35 mL) was added directly to the beads, and total RNA from Input and IP samples was extracted using the RNeasy Micro Kit, including on-column DNase digestion. RNA was eluted in 20 µL RNAse-free water and stored at -80°C. RNA yield and quality were evaluated by Bioanalyzer 2000 (Agilent) using the Total RNA Pico Kit. Before proceeding to library construction, 1 ng of total RNA from each Input and IP sample was used to program a reverse transcription reaction using the Protoscript cDNA synthesis kit (NEB) with random hexamer priming, followed by real-time quantitative PCR (RT-qPCR) on a GeneQuant qPCR machine (BioRad). We performed RT-qPCR for *Esr1*, *Gfap* and *Gapdh* to ensure enrichment or depletion of cell type markers and quality control. For each IP sample we calculated enrichment of *Esr1* and depletion of *Gfap* relative to Input using comparative Ct (Δ/ΔCt). We then used 10 ng of total RNA as input for the Nugen Universal mRNA kit (Tecan) to generate indexed cDNA libraries. Libraries were sequenced on an Illumina HiSeq4000 or NovaSeq6000 to an average depth of ∼32.5 million reads/sample (range ∼16.5-44 million).

### snRNAseq

Esr1+ nuclei from *Esr1^Cre^;SunTag* F_P_ mice were isolated using a modified INTACT protocol as described previously^3,53^. Micro-dissected brain regions were suspended in ice-cold nuclear homogenization buffer (HB) consisting of 0.25 M sucrose, 25 mM KCl, 5 mM MgCl_2_, and 20 mM Tricine-KOH pH 7.8 supplemented with 1 mM DTT, 0.15 mM spermine, 0.5 mM spermidine, EDTA-free cOmplete mini protease inhibitor (Roche), and 40 U/µL murine RNAse inhibitor. The tissue was homogenized, followed by the addition of 60 µL of 5% IGEPAL, and further homogenization. The suspension was filtered through a 40 µm filter and mixed with 1 mL (1 volume) of 50% Optiprep (Sigma-Aldrich), then underlaid with a 30% Optiprep solution and centrifuged at 10,000 *g* for 20 minutes at 4°C to pellet the nuclei. The supernatant was discarded, and the nuclei were resuspended in 1 mL HB supplemented with 1 mM DTT, 40 U/mL RNAse inhibitor, and 0.1% IGEPAL. Nuclear integrity was assessed using Trypan Blue staining and visualization on a hemocytometer. Nuclei were purified using an ARIA FACS. Gates were set by first identifying a side population with 1) high 488 nm (green signal; indicative of *Esr1*/*Sun1*+ nuclei) and low 546 nm (red signal; indicative of auto-fluorescent particles), back-gating to identify events corresponding to nuclei based on side scatter, and further filtering doublets based on forward scatter versus side scatter. Nuclei were sorted into 1 mL of HB supplemented with 1 mM DTT and 40 U/mL RNAse inhibitor, and 0.1% IGEPAL. The yield of sorted nuclei was verified by counting Trypan Blue-stained nuclei on a hemocytometer. The nuclei were pelleted by centrifugation at 2,000 *g* for 10 min at 4°C and resuspended in 60 µL of sterile PBS with 2% bovine serum albumin. They were again inspected and quantified by Trypan Blue staining prior to library preparation using the 10X Genomics Chromium platform with the 10X Genomics RNA 3’ v3.1 kit. Libraries were sequenced on an Illumina NovaSeq6000 to a target depth of 80,000 reads/nucleus to an average depth of ∼267 million reads per library.

### Generation and sequencing of Hi-C libraries

Nuclei from micro-dissected VMHvl of F_R_ and F_P_ *Cckar^Cre/+^;SunTag/+* mice (n=5-6 for each of 2 replicates) were isolated as described above for snRNAseq and fixed with 4% paraformaldehyde (PFA) as described previously^86^. Cckar+ nuclei were purified using FACS as described above for Esr1+ nuclei. Pure nuclei were used for chromosome conformation capture (Hi-C) with the Arima 3C Kit (Arima Genomics) following manufacturer’s instructions. DNA was concentrated using the DNA Clean and Concentrator -5 Kit (Zymo Research). 1 ng of purified DNA/sample was used for library preparation with the DNA Prep, (M) Tagmentation Kit (Illumina). Libraries were sequenced at 150 bp paired-end on an Illumina NovaSeqX Plus to a depth of ∼1.4 billion reads/sample/replicate, totaling ∼5.6 billion reads.

### Bioinformatics

#### TRAPseq analysis and identification of pDEGs

TRAPseq data from F_P_ samples was analyzed as previously described^3^. R1 reads were clipped to 50 bp and processed with bbtools clumpify.sh program (RRID:SCR_016965) to remove exAmp duplicates caused by the NovaSeq6000 patterned flow cell (options: tossbrokenreads dedupe optical spantiles adjacent dupedist = 12500). Remaining reads were pseudoaligned and quantified against mouse polyadenylated transcripts (mm10, Gencode vM21) using Kallisto^87^ with the following options: --bias -b 40 --single -l 180 -s 35. Transcript-length normalized counts were loaded into R using the tximport package (RRID:SCR_016752), and differential expression analysis was performed with DESeq2^88,89^ (RRID:SCR_015687). Mitochondrial genes and genes with fewer than 5 counts in at least 7 libraries were excluded. F_P_ samples were compared to previously generated F_R_ and F_NR_ samples^3^. To check for global batch effects, all F_P_, F_R_ and F_NR_ samples from all four regions were loaded into DESeq2, log_10_-normalized gene expression was calculated separately for each Esr1+ population and physiological state, and principal components analysis (PCA) was conducted using the PCAtools package (RRID:SCR_025593). To identify pDEGs within each region a generalized linear model was constructed for each Esr1+ population using libraries from all three conditions with default DESeq2 parameters. Individual pairwise comparisons (F_P_ v F_R_ and F_P_ v F_NR_) were extracted using the ‘contrast’ function in DESeq2. Genes were considered differentially expressed if they passed a false discovery rate (FDR)-adjusted *p* value <0.05 in at least one pairwise comparison. Unless otherwise noted, for subsequent analyses we only considered genes as pDEGs if they exhibited an absolute fold change ≥1.5. High-fold change (>4-fold) pDEGs were identified using these approaches from TRAPseq datasets. Chromosomal locations of genes encoding pDEGs were obtained using Mouse Genome Informatics Batch Query (https://www.informatics.jax.org/batch). To visualize separation of samples based on DEG, PCAtools was used to plot PC1 and PC2 for each Esr1+ population on log-normalized counts of all pDEGs identified in that population. Separation of samples based on DEGs, PCAtools was used to plot PC1 and PC2 for each Esr1+ population on log-normalized counts of all pDEGs identified in that population.

#### k-means clustering

*k*-means clustering of pDEGs was performed using the base R ‘kmeans’ function. Z-scored expression values for all pDEGs plus previously published estrous-differentially expressed genes (eDEGs)^3^ were calculated across all TRAPseq biological replicates (F_P_, F_R_, F_NR_; n=3 each). This data frame was then subjected to *k*-means clustering with values of *k* ranging from 2 to 10. Optimal cluster number (*k*) was determined as the maximum value of *k* where cluster centroids were consistent within biological replicates of the same condition and region rather than being driven by individual replicates. Once optimal *k* clustering was determined for each population, we annotated similar expression patterns that were evident across regions for visualization and interpretation.

#### Gene Ontology analysis

Gene Ontology (GO) analysis was performed using topGO R package^90^ (RRID:SCR_014798). Enriched GO: Biological Process pathways were identified using a standard pipeline with following options: algorithm = “weight01”, statistic = “fisher”, nodeSize = 5. The union of all pDEGs (F_P_ v F_R_, F_P_ v F_NR_) from each Esr1+ population or alternatively, the union of all pDEGs with >4-fold change (genes considered to be large fold change), served as input, with all expressed genes from that population as background (based on TRAPseq data). Significant GO pathways (*p* < 0.05) were manually categorized. For visualization, we plotted the most significant pathways relevant to female behavior and physiology and pregnancy, shortening redundant pathway names. A complete list of significantly enriched terms can be found in the extended data. Plots were generated using ComplexHeatmap R package^91^ (RRID:SCR_017270). Similarity between significantly enriched GO pathways identified from the entire collection of pDEGs and those identified from pDEGs with >4-fold change was tested using Fisher’s exact test, with all tested GO Biological Process pathways as the background.

#### Disease Ontology analysis

Disease Ontology (DO) analysis was conducted using DOSE R package^92^ (enrichDO function; pvalueCutoff = 0.05, pAdjustMethod = “BH”, minGSSize = 5, maxGSSize = 500, qvalueCutoff = 0.05). Mouse gene symbols were converted to human Entrez IDs using biomaRt R package^93^ (RRID:SCR_019214). The union of all pDEGs from each Esr1+ population or alternatively, the union of all pDEGs with >4-fold change was used as input. Significant diseases (FDR-corrected *p* value <0.05) were manually annotated into categories For visualization, we plotted the diseases more prevalent in females during pregnancy, related to female reproductive tissues, or that are specific to pregnancy. A complete list of significantly enriched terms can be found in the extended data. Redundant disease terms enriched by similar gene sets (e.g., “ovarian cancer” and “ovarian carcinoma”) were omitted. Plots were generated using the ComplexHeatmap R package. Similarity between significantly associated diseases identified from the entire collection of pDEGs and those identified from pDEGs with >4-fold change was tested using Fisher’s exact test, with all diseases included in the enrichDO analysis serving as the background. For the visualization of pDEGs associated with disorders of mood and affect (DOID:3324) and with pre-eclampsia (DOID:10591), we extracted the list of genes from the enrichDO output. Human gene symbols were converted to mouse gene symbols using the biomaRt R package. The average normalized expression levels of these pDEGs across the replicates were *z*-scaled for F_P_, F_R_, and F_NR_, and plotted using the ComplexHeatmap R package.

#### snRNAseq data analysis and cell type identification

Initial quality control analysis and filtering of each library was conducted as previously described^3^. 10X Genomics libraries were pseudoaligned to the mouse transcriptome (mm10, Gencode vM21) using kallisto-bustools^94^. Reads were aligned to a modified transcriptome index that included intronic sequences to quantify unspliced transcripts. Cell x gene matrices were loaded into R using the BUSpaRse package (https://github.com/BUStools/BUSpaRse) and analyzed with Seurat (RRID:SCR_016341). Each F_P_ library from each Esr1+ population was initially analyzed individually for quality control. Cells were retained if at least 500 Unique Molecular Identifiers (UMIs) were detected, while a gene was retained if ≥1 UMI was detected for that gene in ≥3 cells. Count data was variance stabilized in Seurat using the SCTransform function^95^. The abundant non-coding RNA *Malat1* accounted for 1-4% of all UMIs/cell and showed differences in relative abundance between libraries with no systematic correlation with hormonal or reproductive state (not shown). To avoid spurious cell type clustering based on expression of this gene we calculated the percentage of reads mapping to this gene for each cell (% *Malat1*) to use as a regression variable for SCTransform. The top 3000 variable features were identified and used as input for PCA followed by identification of cell types using the Jaccard-Louvain community detection algorithm. Initial clustering was performed using the top 30 PCs and a resolution of 1.5 to identify non-neuronal cell types and cell types consisting of low-quality cells or probable doublets. We identified and removed contaminating non-neuronal cell types (that were always Esr1–) based on enriched expression of the genes *Plp1*, *Mbp*, *Inpp5d*, and *Cxcr1* (enriched in glial, microglial, and endothelial cells). We filtered cell types consisting of low-quality cells or probable doublets based on one or more of the following criteria: uniformly low UMI count (median > 2-fold lower than that of all other cell types), high mitochondrial content (any mitochondrial transcript detected in > 25% of cells), and/or lack of significantly enriched markers relative to all other cells. At each filtering step we removed cell types consisting of low-quality cells, non-neuronal cells, and any cell types in which <25% of cells expressed *Esr1*. Filtering and PCA were iteratively repeated until all remaining cell types were neuronal and Esr1+ in >25% cells.

#### Integration with previously published Esr1+ snRNAseq data

Filtered F_P_ snRNAseq data were integrated with previously published Esr1+ snRNAseq datasets from males and F_R_/F_NR_ females. First, all libraries from all regions and conditions were merged, PCA was performed (top 50 PCs), and nuclei were visualized in a common UMAP space. After confirming nuclei primarily co-embedded by region rather than sequencing batch, sex, or reproductive state, we analyzed each region independently, merging F_P_, F_R_, F_NR_ and male cells without batch correction. For our previous study we identified cell types in each Esr1+ population by performing graph-based clustering in Seurat using the top 30 principal components with the resolution parameter set to 1.2, which were determined by scanning and evaluating multiple combinations of these parameters using the scclusteval package^96^. We elected to analyze the integrated datasets using identical parameters after integrating cells from F_P_. We reasoned that this strategy would allow us to evaluate how robust our previous cellular taxonomy was to the addition of additional samples, identify pregnancy-emergent cell types, and determine if the addition of new cells resulted in any previously annotated cell types being split into finer subdivisions. After re-running PCA and graph-based clustering, we took two approaches to compare our updated (F_P_ inclusive) cell taxonomy to our previously published classification of Esr1+ cell types. First, we extracted the union of all variable genes used for PCA in each dataset (a proxy for cell type marker genes), calculated the average expression of each gene in each cell type using the Seurat AverageExpression function, and generated Peason’s correlation matrix comparing the previous and updated cell types. The resulting correlation matrix was visualized as a heatmap with cell types automatically ordered along a rough diagonal using the slanter package^97^. This approach evaluates whether each previously identified cell type can be matched with corresponding type(s) in the updated taxonomy based on expression of variable genes used for clustering. Second, we modified the scclusteval package to calculate the Jaccard Similarity Index of cell barcodes assigned to each cell type in the previous and updated classification schemes. We then visualized pairwise comparisons of cell types in the two schemes with pheatmap (RRID:SCR_016418) to confirm that cells that were assigned the same cluster in our previous analysis retained their community membership in our new dataset.

#### Further annotation of cell types

We classified cell types expressing *Slc17a6* or *Slc17a7* in >25% of cells as excitatory neurons and those expressing *Gad1* in >25% of cells as inhibitory neurons. To identify genes enriched in individual cell types, we used Model-based Analysis of Single-cell Transcriptomics (MAST; RRID:SCR_016340)^98^, as implemented in the FindAllMarkers function in Seurat with the following parameters: min.pct = 0.25, lgfc.thesh = 0.223, min.diff.pct = 0.25, test.use = “MAST”. This translates into enriched marker genes 1) being expressed in >25% of the cells in the cell type of interest, 2) exhibiting a 25-percentage point difference in number of cells expressing the gene between the cell type and all other cells, and 3) exhibiting ≥1.25-fold difference in log-normalized expression per cell in that cell type versus all other cells. To assign marker gene annotations to cell types, we sought to identify a gene that met these criteria in the cell type of interest and no others within its corresponding Esr1+ population. We annotated cell types lacking such genes based on marker genes enriched in as few other cell types as possible. All marker genes used for annotation were expressed in both TRAPseq and snRNAseq datasets.

#### Mapping pDEGs on to cell types

We defined a gene as ‘expressed’ in a cell type if ≥1 UMI was detected in ≥ 25% of cells in that cell type. To determine whether individual cell types expressed disproportionately more pDEGs in F_P_ or F_R_/F_NR_, we normalized the number of pDEGs upregulated in both conditions for each comparison (F_P_ v F_R_; F_P_ v F_NR_) and then obtained the ratio of the two conditions for that comparison. This ratio for each cell type was scaled across all cell types for that Esr1+ population to obtain a z-score. To determine whether individual pDEGs were significantly enriched or depleted in a given cell type, we calculated z-scored expression of all pDEGs across all cell types in each Esr1+ population and visualized it as a hierarchically clustered heatmap, using the pheatmap package.

#### Identification of condition-biased cell types

To identify condition-biased neuronal states within each Esr1+ population we compared the proportion of cells from each condition (F_R_, F_NR_, and F_P_) assigned to each cell type. A cell type was classified as condition-biased if it met all of following criteria: 1) significant difference in cell counts across conditions (Fisher’s exact test, *p* < 0.05), 2) >3-fold difference in % cells contributing to the cell type between the most- and least-represented conditions, and 3) at least 1% of total cells in the population in the most-represented condition.

#### Generation of transcriptional regulatory networks

We nominated transcription factors (TFs) most likely to govern gene expression differences across cell types, including expression of pDEGs, using the Single-Cell rEgulatory Network Inference and Clustering (SCENIC) algorithm (pySCENIC implementation; RRID:SCR_025802)^54^. As input to the gene regulatory network step, pseudobulk transcriptomes were generated for each cell type by merging snRNAseq data across all four regions. A list of mouse TFs was obtained from pySCENIC (https://github.com/aertslab/pySCENIC/tree/master/notebooks). We manually excluded genes unlikely to be TFs from this list by cross-referencing known DNA-binding domains listed in JASPAR^99^, a curated human TF list^100^, and an independent literature search. Regulatory networks were inferred using TFs expressed in ≥25% of cells in ≥1 cell type in ≥1 region. We retained regulons with normalized enrichment scores ≥2 and included both ‘positive’ (TF + upregulated genes) and ‘negative’ (TF + downregulated genes) regulons. We then extracted the full list of regulons and their predicted target genes from the resulting .loom file using the SCENIC, loomR (https://github.com/mojaveazure/loomR), and AUCell R packages (RRID:SCR_021327)^54,101^. To generate regulatory networks comprised of pDEG-encoded TFs and calculate the distance between nodes in these networks we fed SCENIC regulon data into the igraph R package (RRID:SCR_019225)^102^. TF expression patterns were z-scaled across cell types and visualized as a heatmap using the ComplexHeatmap R package with default hierarchical clustering (Euclidean distance).

#### Hi-C analysis

Bulk Hi-C data was analyzed as previously described^86^, with specific modifications. Briefly, paired-end 150-bp reads were aligned to the mouse genome (mm10, Gencode vM25) using BWA-MEM (-5SP, default parameters)^103^. Chromatin contacts were then generated using hickit (https://github.com/lh3/hickit), by running hickit.js sam2seg, hickit.js chronly, and hickit --dup- dist=1. We visualized chromatin contacts using Juicer Tools (https://github.com/aidenlab/JuicerTools) with parameters “pre -n”. Chromatin A/B compartment values, as quantified by the scA/B metric^86^, of each 1-Mb genomic regions were calculated using “dip-c color2” and then normalized to 0–1 in each sample using MATLAB (“tiedrank”). We compared our chromatin contact maps and scA/B vectors to a previously published dataset of brain cells for neurons^104^ and confirmed that our samples displayed long-range chromatin contacts typical of neurons, although one sample (F_P_ replicate #2) displayed some contacts indicative of astrocytic contamination. For each pDEG set (up- or downregulated pDEGs expressed in #30 Cckar+ and #20 Trim36+ neuronal types), scA/B values were taken from the 1-Mb bin containing each gene’s midpoint based on the mouse genome information and then averaged.

#### Spatial transcriptome analysis

We investigated expression patterns of NPY2R and GAD1 in the human POA using a Seurat Data Object containing 10X Genomics Visium datafile (https://doi.org/10.17863/CAM.111988) from a spatial transcriptomic atlas of the adult human hypothalamus^59^. Slice5A from a female donor (79-yrs age) was selected for visualization because it was the only slice annotated with the POA. We plotted the log10 normalized spatial expression levels of NPY2R and GAD1 using the Seurat ‘SpatialFeaturePlot’ function and hematoxylin-eosin staining for the same slice with the Seurat ‘SpatialDimPlot’ function.

### In situ hybridization

Genes selected for *in situ* hybridization (ISH) validation of TRAPseq data represented: 1) F_P_ vs. F_R_ and F_P_ vs. F_NR_ pDEGs, 2) pDEGs in all four sequenced regions, and 3) a range of fold-changes (1.6 to 11-fold). F_R_ and F_NR_ mice were generated by ovariectomy and hormone priming of C57BL6/J females (8–10 weeks old) exactly as previously described^3^. Mice were deeply anesthetized, transcardially perfused, and brains were fixed overnight in 4% paraformaldehyde at 4°C. Coronal sections (50 µm) were prepared using a vibrating microtome (Leica). Hairpin chain reaction (HCR) ISH was performed exactly as described previously^3^, using probes and fluorescent amplifiers (AlexaFluor-488, 546, or 647) purchased from Molecular Instruments. Up to 40 probe pairs per gene were designed, assigning fluorophores based on gene expression levels (green – high; red – medium; far-red – low) estimated from TRAPseq data. Images were acquired using confocal microscopy (Zeiss LSM800, 20× objective) and processed identically in ImageJ for all pairwise comparisons^105^.

### Caspase mediated ablation and monitoring of pregnancy

Targeted neuronal ablation was performed using caspase-induced apoptosis as previously described^9^. Briefly, virgin female *Npy2r^Cre^*mice (8–14 weeks old) and wild-type (WT) controls received stereotaxic injections (0.3 µL) of a viral mixture (9:1 ratio) containing AAVDJ-EF1α-flex-taCaspase3-2A-TEVp (8×10^12^ vg/mL; UNC Vector Core, AV7165) and AAV2-hSyn-EGFP (2×10^13^ vg/mL; Addgene, 50465-AAV2) into the POA (coordinates: ±0.3 mm ML, +0.5 mm AP, –5.4 mm DV from bregma). Five days post-injection, females were paired with sexually experienced intact males and checked twice daily for vaginal plugs. After confirmation of mating, females were singly housed at 10–17 days after plug (DAP) and weighed daily until delivery or until 20 DAP if no delivery occurred.

### Assessing fertilization

Virgin *Npy2r^Cre^* and WT females injected with a viral vector carrying a Cre-dependent version of caspase-3 (as described above) were monitored for estrous cycle via vaginal cytology from 5 days post virus injection. On the first day of estrus, females were paired with sexually experienced intact males at lights-off and checked every 4 hours for vaginal plugs. At 14 hours post-plug observation, blood glucose levels were measured using the OneTouch Ultra 2 Glucose meter (OneTouch), by collecting a drop of tail blood directly onto a OneTouch Ultra Strip connected to the glucose meter. Oviducts were dissected followed by cardiac puncture and blood collection to measure hormone titers in the serum. Mice were then transcardially perfused, and brains were collected for post-hoc histological analysis. The oviducts were flushed with M2 medium (Cytospring; M2103). Harvested ova were cultured in KSOM based medium (Embryotech; ETECH-EL-50) at 37°C, 5% CO_2_, and assessed after 24 hours. Embryos reaching the 2-cell stage were categorized as fertilized. Ovaries were collected, post-fixed in 4% PFA and stored for subsequent H&E staining.

### Superovulation and embryo collection from donor mice

3-4 weeks old donor C57BL6/J females were superovulated to allow for harvesting of embryos for subsequent embryo transfer as described below. Briefly, donor females received an intraperitoneal injection of 5 IU pregnant mare serum gonadotropin (Aviva; OPPA01037), followed 48 hours later by and intraperitoneal injection of 5 IU human chorionic gonadotropin (Sigma; C8554). Immediately after the second injection, females were paired with sexually experienced intact males. Females were monitored twice daily for vaginal plugs. At 3.5 DAP, blastocysts were collected by flushing uterine horns with M2 medium (Cytospring; M2103) and cultured in KSOM based medium (Embryotech; ETECH-EL-50) at 37°C, 5% CO_2_ until embryo transfer.

### Embryo transfer to recipient mice

Recipient virgin *Npy2r^Cre^* and WT females were injected with viral vector carrying a Cre-dependent version of caspase-3 as described above. Five days after injection, females were paired with sexually experienced vasectomized males and checked twice daily for vaginal plugs. At 2.5 DAP, recipient females received 12–15 blastocysts (E3.5) harvested from donor females into each uterine horn. Females were then singly housed and weighed daily until delivery or until 17 days post-transfer if no delivery occurred.

### Visualizing implantation sites

Implantation sites were visualized through Chicago Blue staining as previously described^69^. Briefly, three days after embryo transfer and blood glucose levels were measured (as described above). Recipient females were anesthetized and blood was retro-orbitally collected. Implantation sites were labelled by injection of 0.1 mL of 1% Chicago Sky Blue 6B dye (Sigma; C8679) into the tail vein. Five minutes post-injection, uteri were dissected, and implantation sites were enumerated based on dye retention. Ovaries were collected, post-fixed in 4% PFA, and stored for subsequent H&E staining. Females were then transcardially perfused, and brains were collected for post-hoc histological validation of caspase-mediated neuronal ablation.

### Ovarian histology

Paraffin embedded microtome sectioning (5 µm) and H&E staining was performed by Histo-Tec Laboratory (Hayward, CA). 10 sections were obtained from each ovary, and follicles and corpora lutea were counted from one randomly selected section for each mouse by an experienced experimenter blinded to relevant variables. We also examined sections preceding and subsequent to the section being analyzed to determine the stage of follicles and corpora lutea.

### Post-mating vaginal cytology assessment

Virgin *Npy2r^Cre^* and WT females injected with Cre-dependent caspase virus (as described above) were paired with sexually experienced vasectomized males five days post-injection. Vaginal plugs were checked twice daily. After plug confirmation, vaginal cytology was checked daily for approximately 20 days to monitor estrous cyclicity^106^. Briefly, mice were immobilized, and the vagina was flushed with 10 μl of saline until liquid became cloudy. Sample was then expelled onto a microscope slide and observed at 20x magnification under bright-field illumination in an epifluorescence microscope (Zeiss Axio Imager A2). Assessment of stage was performed by an experimenter blinded to relevant variables.

### Validation of POA^Npy2r^ neuronal ablation

HCR *in situ* hybridization for *Npy2r* was performed (as described above) to quantify neuronal ablation in all experimental (*Npy2r^Cre^*) and control (WT) females. Brain sections were imaged on a confocal microscope (Zeiss LSM800, 20× objective, 4 optical sections at 7 µm intervals) by an experimenter blinded to relevant variables. *Npy2r*+ cells were quantified using CellProfiler4^107^. Cells were segmented using the IdentifyPrimaryObjects function to segment nuclei based on DAPI staining and defining the soma as occupying a 20-pixel radius beyond the border of the nucleus. ISH signal for *Npy2r* was detected with the IdentifyPrimaryObjects function using the “Global” threshold strategy, the “Robust Background” thresholding method, the “Mean” averaging method, and the “Standard Deviation” variance method. We set an intensity criterion of 2 standard deviations above the mean and a minimum pixel width of 6 and maximum pixel width of 30 to differentiate ISH signal from background. We enumerated Npy2r+ cells in 3 coronal slices per slide through the POA. These cell numbers were plotted as a fraction of cells in the WT/control (# Npy2r+ cells from Npy2r^Cre^ or WT POA ÷ mean # Npy2r+ cells from WT POA for that experiment). Only experimental animals with >70% reduction in *Npy2r*+ cells compared to WT controls were included in the analyses.

### Serum hormone measurements

Blood was collected at the time of euthanasia after deep anesthesia through cardiac puncture, except for the implantation assessment cohort, in which blood was collected retro-orbitally. In this latter cohort, blood collection was performed at 1 PM (end of lights-on period) to coincide with diurnal surge of serum prolactin during early pregnancy^64^. To harvest serum, blood was incubated for 15 min at 37°C, followed by ≥30 min incubation at 4°C. Samples were then centrifuged at 2,700 RCF for 10 min, supernatant was collected and further centrifuged at 5,600 RCF for 10 min to remove any residual red blood cell contamination. Resulting serum was aliquoted and stored at -80°C until analysis. Serum estrogen, progesterone, and prolactin levels were measured using commercial ELISA kits (Cayman 501890, Cayman 582601, Invitrogen EMPRL) following manufacturer’s protocol. Serum FSH and LH were measured by the University of Virginia Ligand Assay & Analysis Core using custom ELISAs^108,109^.

### Statistics and reproducibility

Bioinformatics analyses were performed using R and Python, with statistical tests and criteria detailed in the relevant method sections. Functional studies were analyzed blind to relevant variables including genotype or experimental condition, and statistical analyses were performed using GraphPad Prism (RRID:SCR_002798). Categorical data were analyzed by Fisher’s exact test. Non-categorical data were assessed for outliers using the ROUT method (Q=1%) and tested for normality using the D’Agostino-Pearson omnibus test. Two-group comparisons were performed using *t*-test (parametric) or Mann-Whitney test (non-parametric). Multiple comparisons were performed using two-way ANOVA followed by Sidak multiple comparison test. Weight gain data were analyzed using repeated-measures ANOVA (parametric). The correlation between the number of *Npy2r*+ cells and the number of zygotes was assessed using Pearson’s correlation coefficient. Sample sizes for each experiment are indicated in the text or figure legends.

## Author contributions

ST, JRK, VMAC, and NMS conceived and designed the study. JRK performed TRAPseq and snRNAseq experiments, with help from AT. ST performed Hi-C experiments, with help from JRK and advice from BP and LT. ST, JRK, MW, and LT performed analysis of all sequencing studies. ST, JRK, VMAC, MA, AT, SAEP, and COD performed histology or helped with histological analyses. ST, VMAC, MA, and YW performed embryo transfers, fertilization check, vaginal cytology, or hormone titer measurements. ST, JRK, YW, RY, and NMS performed dissections for all sequencing studies. MHF and DJL provided advice on handling early embryos, superovulation, and ovarian histology and analysis. ST, JRK, VMAC, and LC maintained the mouse colony. ST, JRK, VMAC, and NMS conceptualized data visualization and prepared Figures. ST, JRK, VMAC, and NMS wrote the manuscript with input from all authors.

---

**Extended Data Figure 1:**
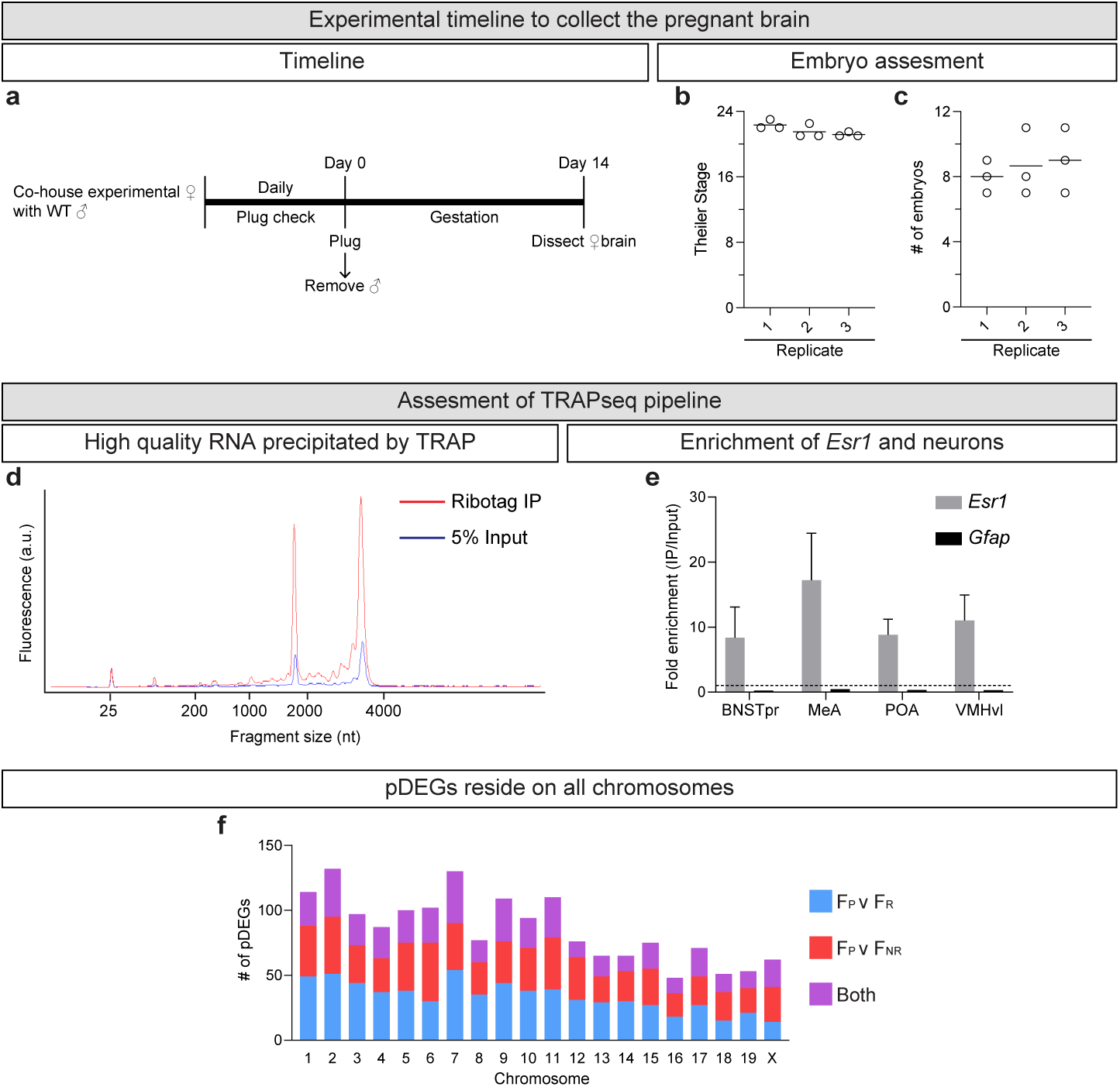
TRAPseq quality control. **a.** Schematic of timeline to generate F_P_ mice. **b-c.** Theiler Stage of embryos (**b**) and number of embryos (**c**) from F_P_ mice used for TRAPseq. Each hollow circle represents one F_P_ mouse. Horizontal lines denote Mean. Replicate, biological replicate. **d.** Representative Bioanalyzer traces showing high-quality RNA precipitated from Esr1+ neurons. Red, immunoprecipitated RNA; blue, pre-immunoprecipitation RNA (‘Input’). **e.** Real-time quantitative PCR analysis following RNA immunoprecipitation demonstrates enrichment of *Esr1* mRNA (gray) and depletion of the astrocytic marker gene *Gfap* mRNA (black). Dashed line = 1. Mean ± S.E.M*. N* = 3 biological replicates. **f.** Distribution of pDEGs across chromosomes. Blue, pDEGs identified in the F_P_ v F_R_ comparison; red, pDEGs identified in the F_P_ v F_NR_ comparison; purple, pDEGs identified in both F_P_ v F_R_ and F_P_ v F_NR_ comparisons.

**Extended Data Figure 2:**
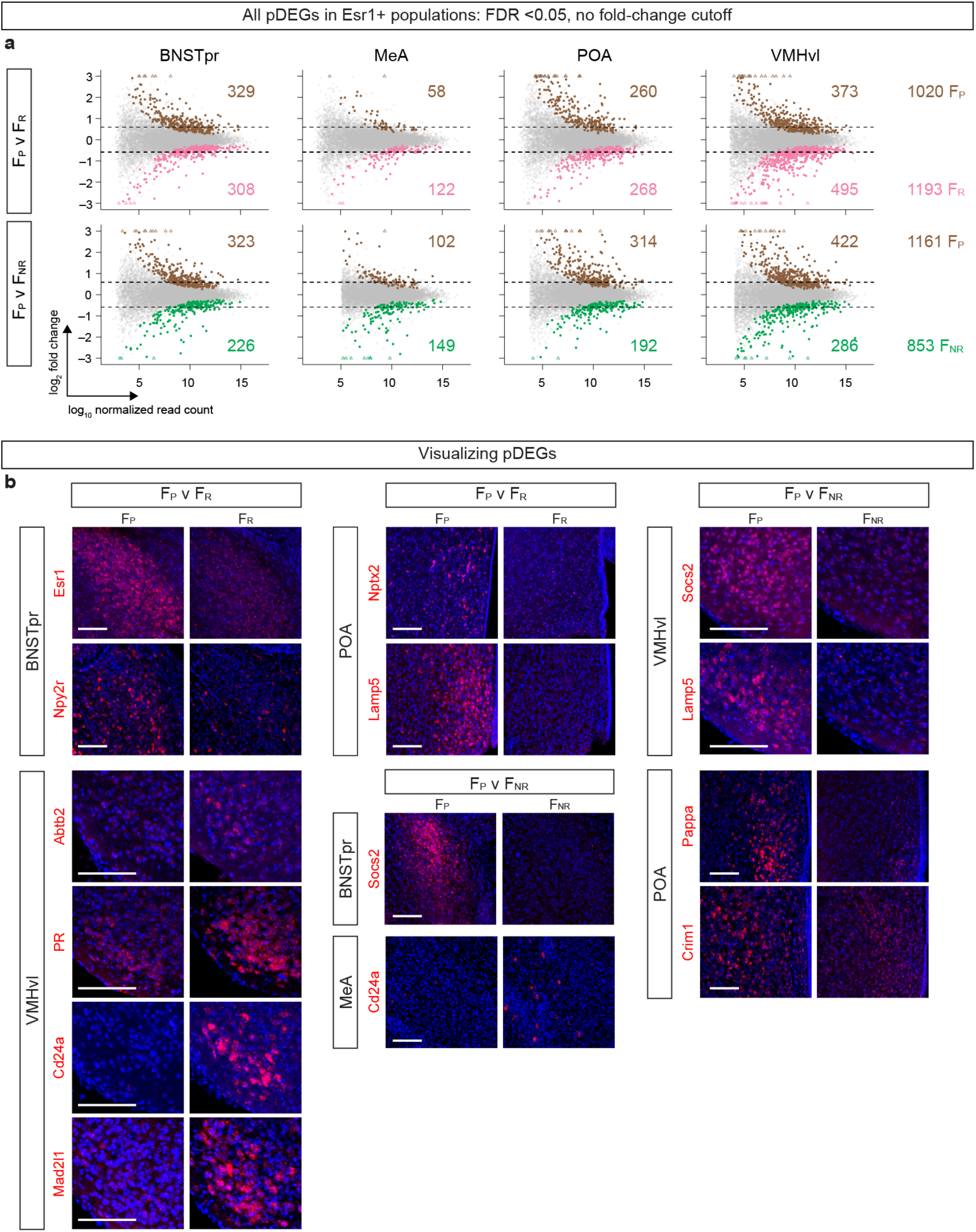
Further characterization and validation of pDEGs. **a.** Scatter plots of all genes that pass FDR-adjusted *p* value < 0.05 regardless of fold-change in expression difference. Dots represent individual genes with FDR-adjusted *p* value < 0.05 (significant; colored) or > 0.05 (not significant; gray) between comparisons. Triangles represent genes whose expression changes exceed the scale. Dashed black lines denote a 1.5-fold differential expression threshold. Colored numbers enumerate genes significantly upregulated in each condition and comparison or (far right) across all regions combined. *N* = 3/condition. **b.** Representative images of ISH for pDEGs. Red, mRNA; blue, DAPI. Scale bars, 100 µm. *N* = 2/condition/gene.

**Extended Data Figure 3:**
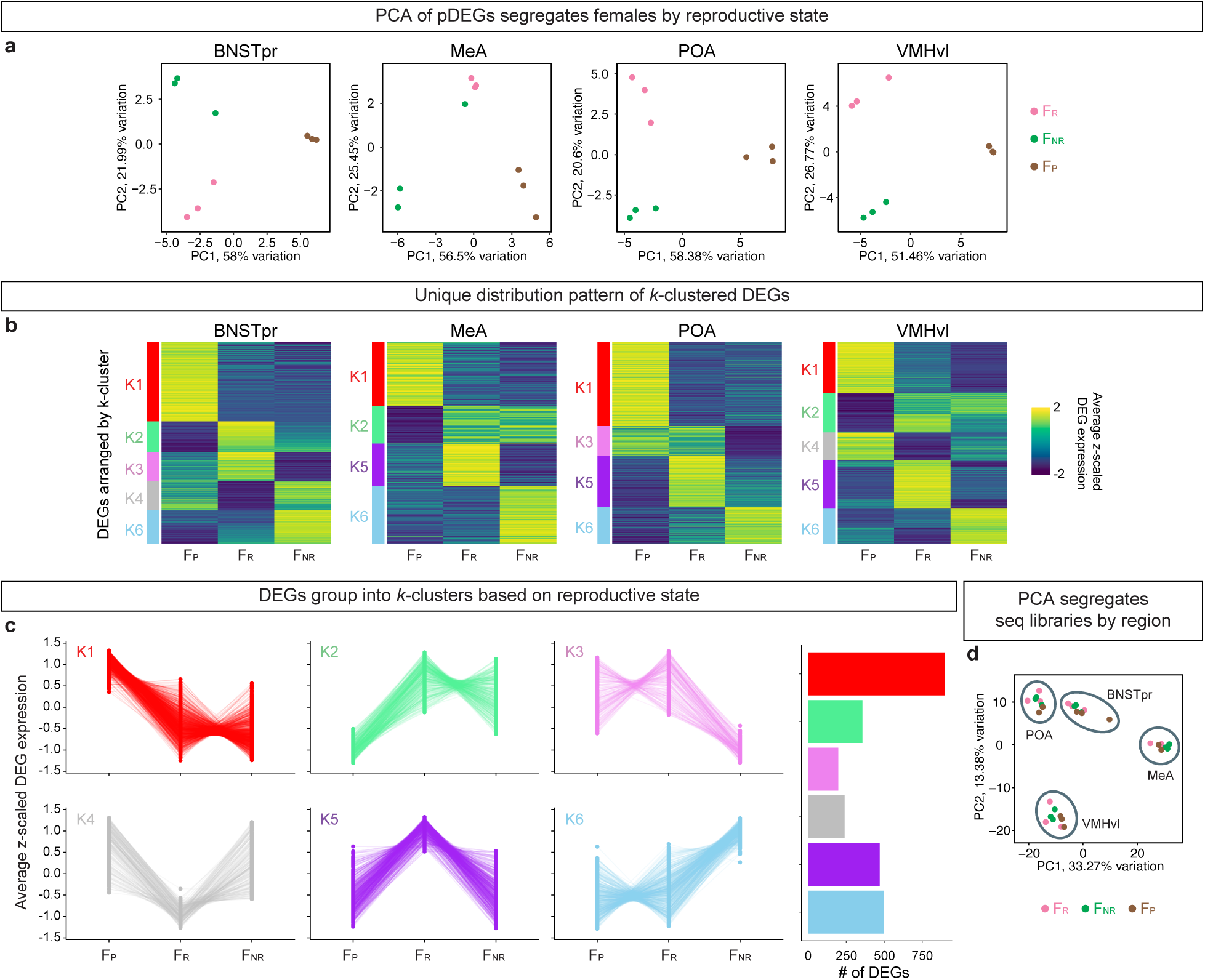
Molecular signatures of pDEGs. **a.** PCA of pDEGs shows that Esr1+ populations from F_P_, F_R_, and F_NR_ occupy distinct regions in PCA space. **b.** Heatmap of average *z*-scaled expression of pDEGs and previously identified DEGs^3^ between F_R_ v F_NR_, arranged by *k*-means clustering. Differently colored bars on left of each heatmap indicate qualitatively similar *k*-clusters independently identified within each Esr1+ population. **c.** Silhouette plots showing *z*-scaled expression of DEGs within each *k*-cluster across reproductive states (left); total number of DEGs assigned to each *k*-cluster (right). **d.** PCA of pDEGs shows that Esr1+ neuronal populations cluster by region rather than reproductive state.

**Extended Data Figure 4:**
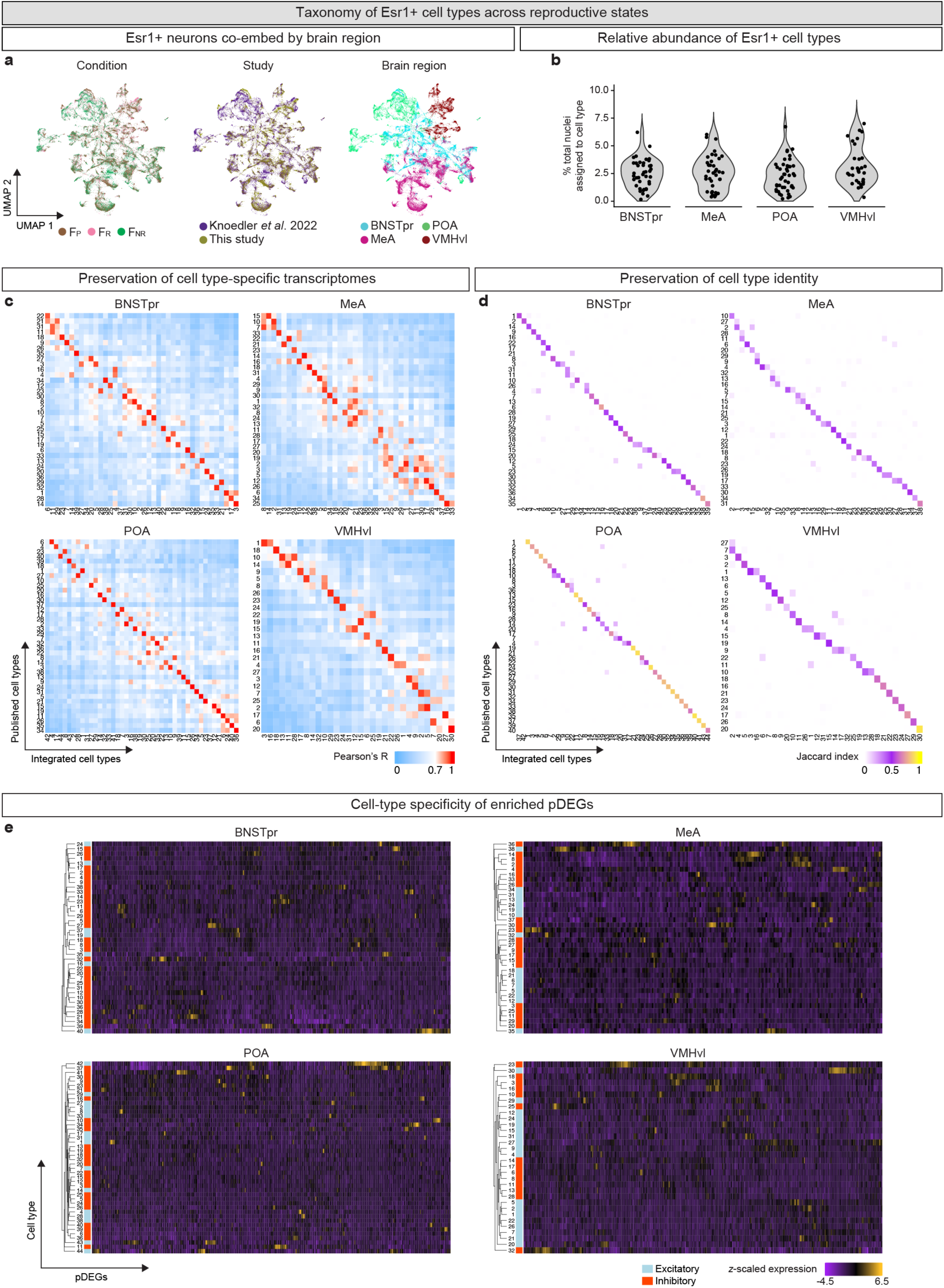
Characterization of transcriptomically-defined neuronal cell types. **a.** UMAP plots demonstrating that Esr1+ nuclei from F_P_, F_R_, and F_NR_^3^ co-embed by region (right) rather than reproductive state (left) or experimental batch (center). **b.** Distribution of relative abundance of Esr1+ neuronal types in each region. **c.** Pearson’s correlation coefficients comparing gene expression signatures between previously published neuronal types^3^ (y-axis) and newly integrated neuronal types from this study (x-axis). High correlations indicate preservation of neuronal type taxonomies. **d.** Jaccard similarity index-based comparison of individual barcoded snRNAseq libraries between previously published neuronal types^3^ (y-axis) and newly integrated neuronal types from this study (x-axis). High similarity index indicates that neuronal type groupings are preserved following the addition of F_P_ nuclei. **e.** Heatmap of *z*-scaled expression of pDEGs showing that particular pDEGs exhibit enrichment or depletion within specific excitatory or inhibitory neuronal types for all Esr1+ populations.

**Extended Data Figure 5:**
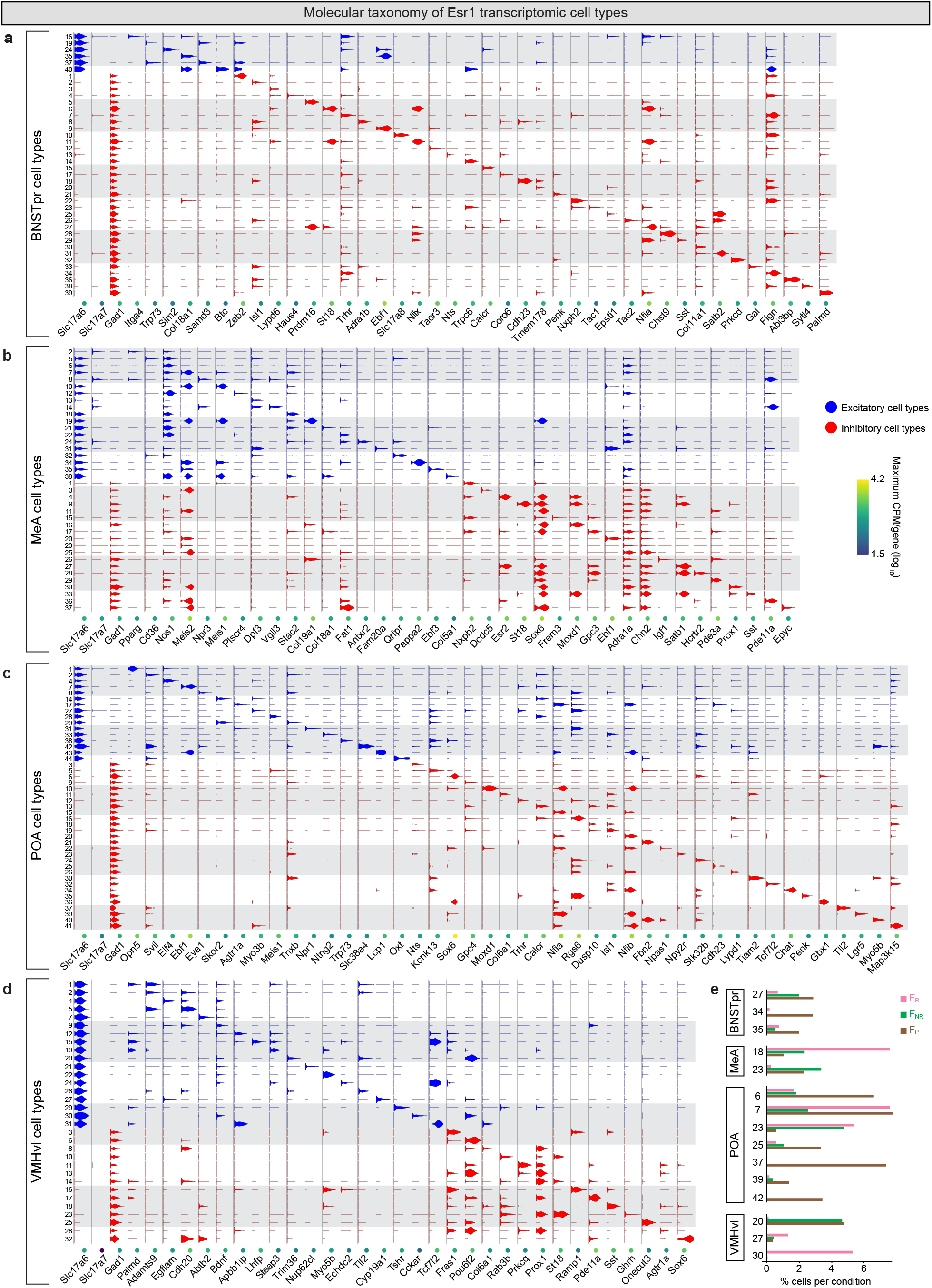
Molecular identity of transcriptomically-defined neuronal cell types. **a-d**. Classification of Esr1+ neuronal types (rows) according to fast neurotransmitter categories (excitatory or inhibitory) and enriched marker genes in the BNSTpr (**a**), MeA (**b**), POA (**c**), and VMHvl (**d**). CPM, counts per million. Sample sizes: *N* = 16,454 nuclei (BNSTpr), 27,713 nuclei (MeA), 20,383 nuclei (POA), and 9,823 nuclei (VMHvl). **e.** Neuronal cell types showing enrichment or depletion in particular reproductive states.

**Extended Data Figure 6:**
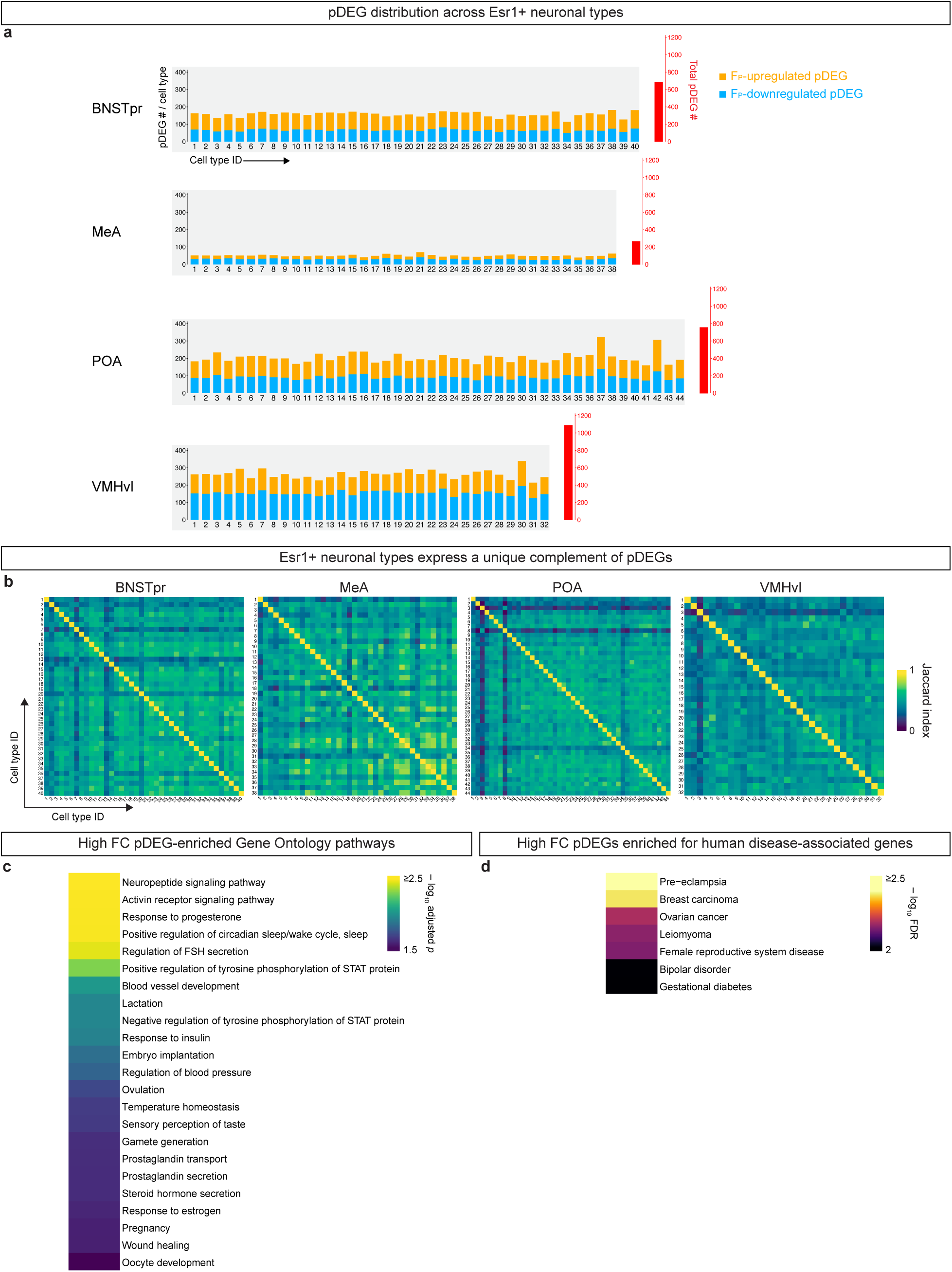
Neuronal type and molecular signatures of pDEGs. **a.** Each neuronal type in all Esr1+ populations expresses a subset of pDEGs in that region. **b.** Each neuronal type in all Esr1+ populations expresses a unique complement of pDEGs. **c.** Gene Ontology Biological Process pathways enriched among the combined set of high fold-change pDEGs (fold-change >4; merged from all Esr1+ populations). Significantly enriched GO pathways (adjusted *p*-value <0.05) related to female biology or pregnancy are shown. **d.** Human diseases significantly associated with the combined set of high fold-change pDEGs (fold-change >4; merged from all Esr1+ populations). Significantly associated diseases (FDR-adjusted *p*-value <0.05) related to female biology or pregnancy are shown.

**Extended Data Figure 7:**
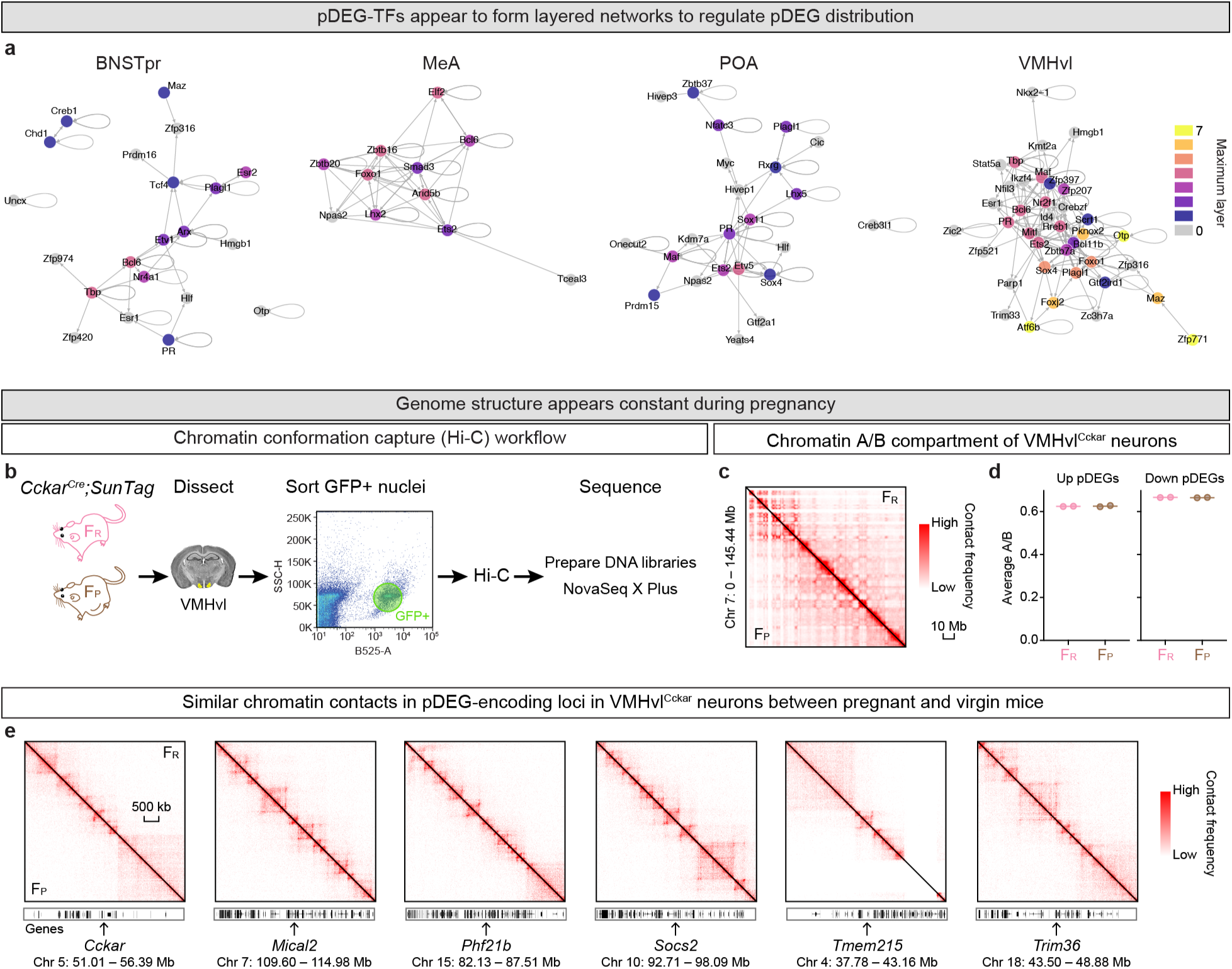
Potential regulatory mechanisms of pDEGs. **a.** Hierarchical regulatory network of pDEG-TFs. Colors indicate the maximum number of steps linking each pDEG-TF to its predicted pDEG-TF targets. **b.** Hi-C workflow for VMHvl^Cckar^ cells. **c.** Chromatin contact map of chromosome 7 of VMHvl^Cckar^ cells showing Hi-C data generated from the limiting number of cells from F_P_ and F_R_ mice. ‘Low’ on the heatmap color scale indicates 0 (no contacts detected between loci), while ‘high’ is proportional to each sample’s total contact number, 938 and 845 contacts/Mb^2^ for F_R_ and F_P_, respectively. N = 2/condition. **d.** No difference in average chromatin A/B compartment residency for pDEGs upregulated (Up) or downregulated (Down) in F_P_ in VMHvl^Cckar^ cells. **e.** No difference in chromatin contacts for representative pDEG-encoding loci in VMHvl^Cckar^ cells between F_P_ and F_R_ states. ‘Low’ on the heatmap color scale indicates 0, while ‘high’ is chosen to be proportional to each sample’s total contact number, 7.3 and 6.6 contacts /10 kb^2^ for F_R_ and F_P_, respectively.

**Extended Data Figure 8:**
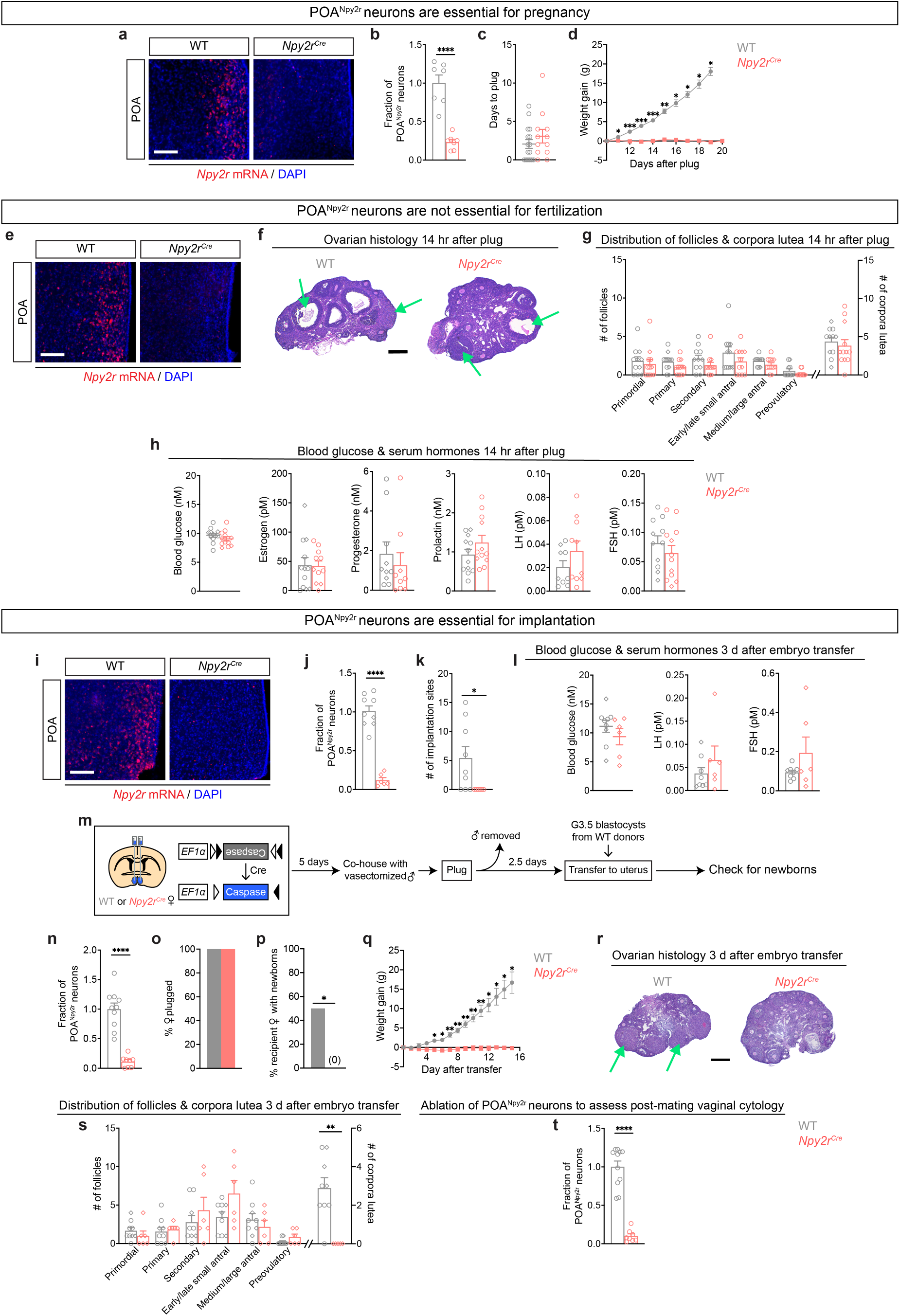
POA^Npy2r^ neurons are essential for viable pregnancy. **a-d.** Ablation of POA^Npy2r^ neurons (**a**, **b**) does not impede successful mating (**c**), but it eliminates the expected gain in weight (**d**). POA^Npy2r^ neuron number plotted as fraction of neurons compared to control for all panels quantifying ablation of these neurons. Scale bar = 100 µm. **e-h.** Ablation of POA^Npy2r^ neurons (**e**). No difference in ovarian histology for follicles and corpora lutea (**f, g**) (**f,** green arrows show different stages of corpora lutea) or blood glucose and a battery of reproductive hormone titers (**h**) 14 hours after successful mating (vaginal plug confirmed) between females ± POA^Npy2r^ neurons. Circles and diamonds, females with and without eggs in oviducts, respectively. LH, luteinizing hormone; FSH, follicle stimulating hormone; scale bar (**e, f**) = 100 µm. **i-l.** Ablation of POA^Npy2r^ neurons (**i**). Three days after blastocyst transfer, females lacking POA^Npy2r^ neurons (**i, j**) have no implanted embryos (**k**) and they have unaffected titers of glucose, LH, and FSH (**l**). Scale bar = 100 µm. **m-q**. Workflow to test role of POA^Npy2r^ neurons in viable gestation after WT blastocyst transfer (**m**). Ablation of POA^Npy2r^ neurons (**n**) does not impede successful mating (**o**) but abrogates viable gestation (**p**) and gain in weight at any point after blastocyst transfer (**q**). **r-s**. Three days after blastocyst transfer, ovaries from females lacking POA^Npy2r^ neurons do not have corpora lutea (**r,** green arrows point to mature corpora lutea in WT females). Circles and diamonds indicate females with and without implantation sites, respectively. Scale bar = 100 µm. **t**. Successful ablation of POA^Npy2r^ neurons for experiments designed to test vaginal cytology. Sample size: WT, *N* **=** 12 and *Npy2r^Cre^*, *N* = 10 (**a-d**); WT, *N* = 12 and *Npy2r^Cre^*, *N* = 12 (**e-h**); WT, *N* = 10 and *Npy2r^Cre^*, *N* = 8 (**n-q**); WT, *N* = 9 and *Npy2r^Cre^*, *N* = 6 (**i-l**, **r-s**); WT, *N* = 11 and *Npy2r^Cre^*, *N* = 8 (**t**). Statistical analyses: Student’s *t*-test (**b**, **g** [# of corpora lutea], **h** [except FSH], **j, n**); Mann-Whitney test (**c**, **h** [FSH], **k-l, s** [# of corpora lutea], **t**); Fisher’s exact test (**o-p**); repeated-measures ANOVA followed by Sidak multiple comparison test (**d**, **q**); two-way ANOVA followed by Sidak multiple comparison test (**g** [# of follicles], **s** [# of follicles]). *, *p* < 0.05; **, *p* < 0.01; ***, *p* < 0.001; ****, *p* < 0.0001.

